# NeuroMechFly v2, simulating embodied sensorimotor control in adult *Drosophila*

**DOI:** 10.1101/2023.09.18.556649

**Authors:** Sibo Wang-Chen, Victor Alfred Stimpfling, Thomas Ka Chung Lam, Pembe Gizem Özdil, Louise Genoud, Femke Hurtak, Pavan Ramdya

## Abstract

Discovering principles underlying the control of animal behavior requires a tight dialogue between experiments and neuromechanical models. Until now, such models, including NeuroMechFly for the adult fly, *Drosophila melanogaster*, have primarily been used to investigate motor control. Far less studied with realistic body models is how the brain and motor systems work together to perform hierarchical sensorimotor control. Here we present NeuroMechFly v2, a framework that expands *Drosophila* neuromechanical modeling by enabling visual and olfactory sensing, ascending motor feedback, and complex terrains that can be navigated using leg adhesion. We illustrate its capabilities by first constructing biologically inspired locomotor controllers that use ascending motor feedback to perform path integration and head stabilization. Then, we add visual and olfactory sensing to this controller and train it using reinforcement learning to perform a multimodal navigation task in closed loop. Finally, we illustrate more biorealistic modeling in two ways: our model navigates a complex odor plume using a *Drosophila* odor taxis strategy, and it uses a connectome-constrained visual system network to follow another simulated fly. With this framework, NeuroMechFly can be used to accelerate the discovery of explanatory models of the nervous system and to develop machine learning-based controllers for autonomous artificial agents and robots.

## Introduction

How the nervous system controls behavior is a deeply entangled problem: nested feedback loops occur at multiple levels including the physiology of individual neurons, the recurrent dynamics of neural circuits, biomechanical interactions with the environment, and sensory signals resulting from one’s own actions. Therefore, investigating and modeling any of these elements in isolation, although useful, will always be fundamentally incomplete. To overcome this gap, neuroscience requires simulation frameworks that enable the exploration of hierarchical feedback loops in an end-to-end manner. Numerous detailed neuromechanical models have been developed to explore how animals control motor programs like walking [1, 2], swimming [3], and transitions between them [4]. As well, in the field of reinforcement learning more abstract and simplified “creatures” [5] have been widely used to model visuomotor coordination [6], decision making [7], and learning algorithms [8]. These latter models often lack realistic motor control and biomechanics and, as a consequence, typically generate only simplified control signals that drive unrealistically limited categorical variables or joint degrees of freedom [9]. Thus, although progress is being made [10–15], a significant gap remains at the interface between machinelearning models and morphologically realistic neuromechanical models of most animals. An ideal model would enable the exploration of *hierarchical* controllers [16]—like those found in biological agents—which include higher-order brain-like systems that integrate multi-modal sensory inputs, ascending motor feedback, and internal states as well as lower-level motor systems that decode descending brain commands and execute behaviors **(Figure 1a)**.

**Figure 1:**
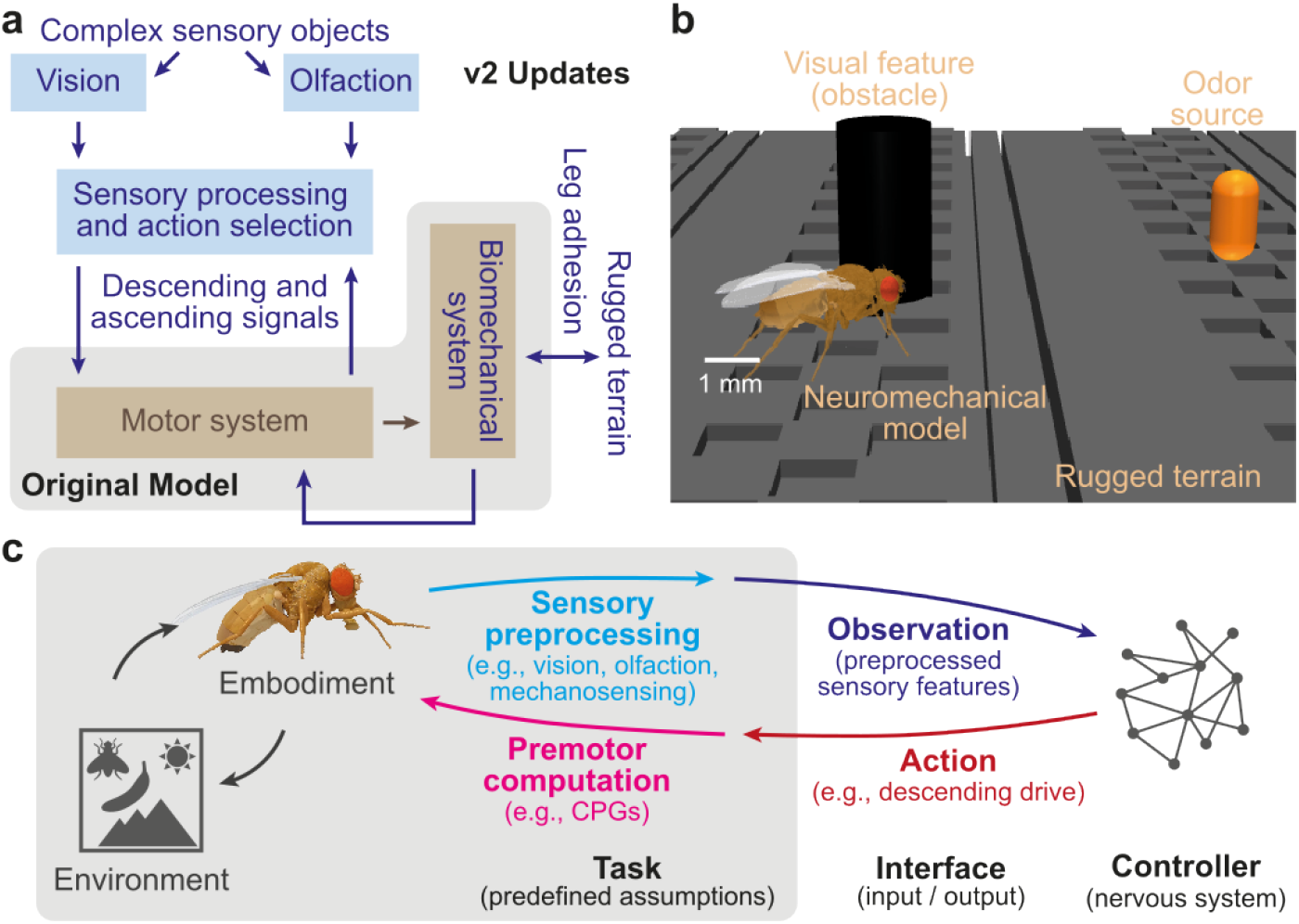
Schematic overview of the NeuroMechFly v2 modeling framework. **(a)** Diagram highlighting the integrative and hierarchical aspects of the expanded NeuroMechFly framework. Sensory inputs—visual and olfactory—are processed by the brain to select an appropriate action. Instructions to execute the selected action are then communicated to the lower-level motor centers (Ventral Nerve Cord or VNC) through descending signals. The VNC executes the motor program via joint actions and mechanosensory feedback signals are sent back to the motor system to inform the next movement and to the brain via ascending pathways. Indicated are components that were present in the original model but improved (brown, in shaded box) as well as new elements (blue, outside shaded box). **(b)** Simulation camera view of the neuromechanical model, NeuroMechFly, walking over complex terrain, using vision to avoid a pillar, and using olfaction to reach an attractive odor source (orange). **(c)** The biomechanical model and its interaction with the environment are encapsulated as a Partially Observable Markov Decision Process (POMDP) task. A user-defined controller interfaces with the task through actions (red) and observations (dark blue). The user can extend the POMDP task by adding preprogrammed processing routines for sensory inputs (light blue) and premotor outputs (magenta), to modify the action and observation spaces handled by the controller.

The adult fruit fly, *Drosophila melanogaster*, is an ideal animal for comprehensively modeling hierarchical control. It has only ∼200,000 neurons in its brain [17] (about 10^3^–10^6^ times smaller than the mouse and human brains); its principal motor system (the ventral nerve cord or VNC) has only ∼15,000 neurons [18] (about 10^2^–10^4^ times smaller than the mouse and human spinal cords). Using this small number of neurons, flies can nevertheless generate complex behaviors. They can walk over complex terrain [19], make rapid corrective maneuvers during flight [20], execute courtship sequences [21], fight competitors [22], and learn [23]. The organization of the fly’s nervous system resembles that of vertebrates yet it has been completely mapped in connectomes (i.e., wiring diagrams) of the brain [24] and VNC [25, 26]. Additionally, thousands of transgenic fly lines enable the repeated targeting of sparse sets of neurons [27] for optical activation [28], silencing [29], and recordings [30]. These resources and tools have led to the development of connectome-constrained models of neural circuits [31, 32]. However, we still lack an integrative simulation framework of the fly, embedded in a physics simulator, to embody and explore these models to identify the principles governing biological intelligence and autonomous behavioral control.

Towards this goal, we previously developed NeuroMechFly, the first morphologically realistic model of the adult fly [13]. With its biomechanical hull, we could infer unmeasured forces and collisions during the simulated replay of recorded limb kinematics from walking and grooming flies. Furthermore, we could use simple coupled oscillators to control fast and stable tethered walking. Although foundational, this previous simulation framework could not fully model hierarchical sensorimotor control: complex environments, brain-level sensory processing, and the physics and biomechanics required for untethered behavioral control were lacking. Here we describe NeuroMechFly v2, a simulation framework that addresses these gaps by (i) improving biomechanics and stepping, (ii) adding leg adhesion, (iii) simulating visual and olfactory sensing, and (iv) enriching the fly’s environment with rugged terrain, obstacles, and sensory objects including other flies **(Figure 1a-b)**. We illustrate the exploration of locomotor control over rugged terrain, simple visual object tracking, and simple odor taxis. Next, we demonstrate the use of ascending limb motor signals to perform path integration and head stabilization. Then, we combine these elements to build integrated hierarchical machine learning models that solve a multimodal task—visually avoiding an obstacle to reach an attractive odor source over rugged terrain. We show how this artificial neural network controller can be trained using reinforcement learning. Finally, we illustrate more biological realism by modeling a *Drosophila* odor taxis strategy to navigate a complex odor plume and using a connectome-constrained visual system network to perform fly following. The modularity of NeuroMechFly v2 allows users to flexibly interact with the simulation at multiple levels of abstraction and facilitates its widespread adoption for research and education. Our implementation’s compliance with a standard reinforcement learning task interface can also facilitate a dialogue between neuroscience, machine learning, and robotics **(Figure 1c, Supplementary Note 1)**.

## Results

### The FlyGym package: a standardized simulation framework

To improve the usability of the NeuroMechFly simulation framework (released as the FlyGym Python package, https://neuromechfly.org/), we made three fundamental changes. First, the package fully complies with Gymnasium [5], a standard interface for controller-environment interaction in robotics and reinforcement learning **(Figure 1c)**. Second, we moved the simulation framework from PyBullet to MuJoCo [33], a more intensively maintained and widely used physics simulator. MuJoCo is known for better stability and performance [34] and supports a wider range of actuators including those for leg adhesion. It was made open-source after NeuroMechFly v1 was published. Third, to facilitate implementing custom environmental features within the simulation, we expanded the interface for the fly model’s arena.

In NeuroMechFly v2 we clarify the framing of the control problem as a Partially Observable Markov Decision Process (POMDP). At each time step, the simulation provides the controller with an observation and, optionally, a user-defined reward. Then, the simulation receives an action from the controller and steps the physics forward accordingly. The observation space is a user-configurable subset of the state space including visual input, olfactory input, ground contacts, joint states (angles, angular velocities, and torques), and the states (e.g., position, orientation) of potentially multiple fly models in the arena. The action space includes the control signals (e.g., target angles) for each actuated degree of freedom (DoF) and the on/off signal for leg adhesion. Users can easily extend this framework by incorporating additional sensory or premotor processing into the Markov Decision Process **(Figure 1c)**. For example, in a visual taxis example described below, we programmed the centroid calculation to reduce the observation space to fewer dimensions (e.g., the azimuth of the object seen from each eye) and used a network of central coupled oscillators to reduce the action space to a twodimensional descending command. This increased flexibility and usability allows one to rapidly adapt the simulation to their own research question.

### Improved morphological accuracy and kinematic realism

A promise of biomechanical models such as NeuroMechFly is to allow researchers studying body movements to, for example, infer unmeasured collisions, contacts, and forces during the replay of real recorded body part kinematics [13]. However, doing so at a high precision is only possible with a morphologically realistically rigged body model. In particular, modeling behaviors that depend on self-contact (such as grooming) requires precise kinematic replay to read out where and how individual body parts interact with one another. Therefore, we improved the morphological accuracy and granularity of several body part meshes. First, we adjusted the placement and default angles of joints between the thorax and front leg coxae as well as between the thorax and head **(Extended Data Figure 1a–b)**. These adjustments were made based on high-magnification video data. Second, to better facilitate control of the antennae and readout of their mechanosensory signals, we split their meshes into three segments: pedicel, funiculus, and arista. We added degrees of freedom (DoFs) between these segments, allowing each to be separately actuated or passively moved and its angular displacement to be measured **(Extended Data Figure 1c)**. We note that users can simplify body geometries if morphological accuracy is of less importance than computational speed.

We also improved the realism of limb kinematics during walking. Leg kinematics in NeuroMechFly v1 [13] were based on data from tethered flies walking on a spherical treadmill. In our simulation these kinematics appeared unnatural during untethered walking, turning, and locomotion over rugged terrain. Therefore, to obtain realistic untethered 3D leg kinematics, we designed a system with three views of a fly walking straight through a corridor. We annotated and triangulated keypoints from walking fly data and extracted individual steps. For each pair of legs we segmented and processed a step while enforcing symmetry, closure, and equal lengths **(Extended Data Figure 2; Video 1)**. Replaying and looping these steps in NeuroMechFly **(Video 2)** drove straight walking more closely resembling that of a real fly.

### Leg tip adhesion enables locomotion in three dimensions

Insects, including flies, use highly specialized adhesive structures to walk over complex 3D terrain with ease. These include adhesive pads with substantial normal forces (*>*100 times body weight) and frictional forces [35]. Adhesion provides mechanical coupling between the legs during locomotion and improves force transduction with the ground. We cannot easily model the physics of real adhesion. Therefore, we modeled leg adhesion as an additional normal force when a leg tip (i.e., pretarsus) is in contact with the surface **(Figure 2a, Video 3)**. As for insects [35], this normal force also increases the frictional forces. Despite the huge forces generated by adhesive pads, insects appear to be able to lift their legs without much effort. Although liftoff mechanisms are known for some insects [36, 37] they are not known for *Drosophila melanogaster*. Therefore, we abstracted the mechanisms used by other insects and lift the legs during walking by turning off adhesion forces during the swing phase. To illustrate how leg adhesion expands the behavioral repertoire of our model, we simulated tripod-gait walking [38, 39] over terrain with up to 180° of inclination**(Figure 2b, Video 4)**. Without adhesion, the fly slipped at as low as 30° inclination. By contrast, with adhesion the fly could locomote over terrain with sometimes more than *>* 90° of inclination **(Figure 2c**). We expect experimental recordings of real inverted walking kinematics to enable locomotion at even higher inclinations.

**Figure 2:**
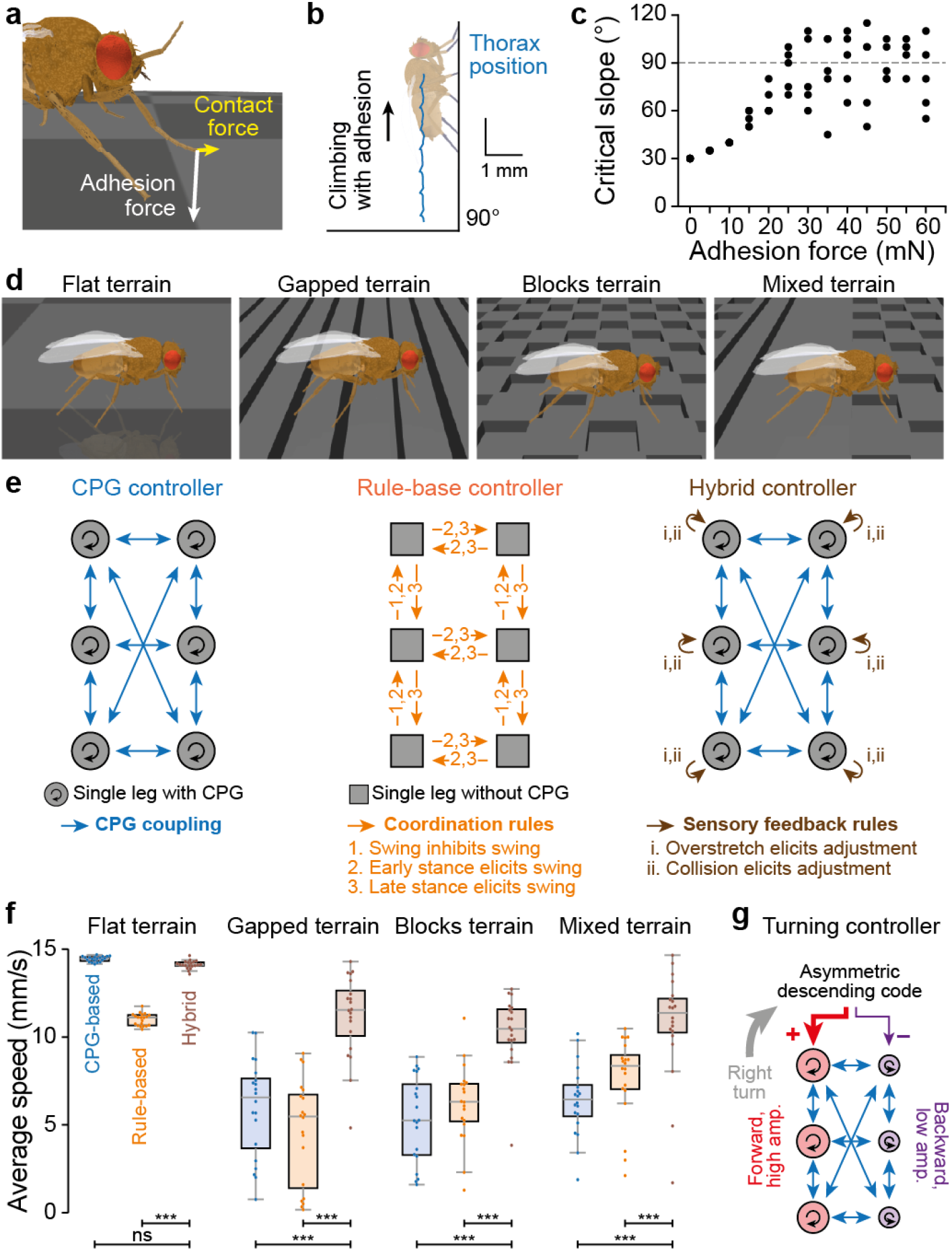
Locomotion in three dimensions and over rugged terrains. **(a)** A schematic of contact forces on the fly’s leg with the addition of adhesion. **(b)** Trajectory (blue) of the fly as it walks up a vertical wall. The legs are controlled by a CPG controller with adhesion enabled (F = 40 mN). **(c)** Critical slope (angle) at which the fly falls or does not proceed forward at different magnitudes of leg adhesion force. **(d)** Terrains for exploring the efficacy of distinct locomotor strategies. We tested four terrains: a flat surface, a surface with gaps, a surface with blocks, and a mixture of all surface types. **(e)** Three controllers tested across terrains: a controller with six coupled CPGs controlling the swing and stance of the legs, a rule-based controller in which the phase of one leg’s movements influences the movements of neighboring legs, and a hybrid controller consisting of coupled CPGs with sensory feedback-based corrective mechanisms that execute stepping phase-dependent adjustments when the leg might be stuck. In all cases leg adhesion is present. **(f)** The performance (average speed) of each locomotor controller while walking over four types of terrain. Shown as dots are *N* = 20 trials, each with a random spawn location and controller initialization. Overlaid are box plots indicating the median, upper and lower quartiles, and whiskers extending to the furthest points excluding outliers that are more than 1.5 *×* the interquartile range (IQR) beyond the IQR. One-sided, asymptotic Mann– Whitney *U* test was used to generate the statistics: “ns” not significant; ** *p <* 0.01, *** *p <* 0.001 (see **Supplementary Table 1** for complete statistics). **(g)** Turning is controlled by the asymmetric modulation of a two-dimensional descending command signal that regulates the directions and amplitudes of oscillators on each side of the body.

### Complex terrains demonstrate the use of locomotor feedback

A variety of mechanisms have been proposed for insect locomotion ranging across a spectrum from those depending purely on central pattern generators (CPGs, namely circuits in the *central* nervous system that produce rhythmic motor output without rhythmic input [40]) to those relying on sensory feedback-based rules [1]. Evidence for each of these control strategies has been found across species, motivating their application in robotics [40–42]. While walking over flat terrain can be solved using a variety of feedback-independent control strategies, leg mechanosensory signals are thought to be required to navigate rugged terrain. To demonstrate that NeuroMechFly can serve as a test bed to evaluate different control strategies in complex environments, we developed three rugged terrain types to compare with smooth terrain **(Figure 2d, “Flat terrain”)**: one with gaps perpendicular to the initial heading of the fly **(Figure 2d, “Gapped terrain”)**, one with blocks of alternating height **(Figure 2d, “Blocks terrain”)**, and one that is a mixture of the previous two **(Figure 2d, “Mixed terrain”)**.

We next built controllers and benchmarked them over flat and rugged terrains. The control strategies tested include purely CPG-based **(Figure 2e, “CPG controller”, Video 5)**, purely sensory feedback rule-based **(Figure 2e, “Rule-based controller”, Video 6)**, or intermediate to these two, with CPGs but also sensory feedback rules to recover from challenging positions **(Figure 2e, “Hybrid controller”, Video 7)**. At baseline (i.e., on flat terrain), CPG and hybrid controllers were fastest **(Figure 2f, blue)**. However, on rugged terrain the CPG-based controller struggled compared with the rule-based controller **(Figure 2f, orange)**. The hybrid controller leveraging both CPGs and sensory feedback rules overcame this trade-off: it remained fast over rugged terrain **(Figure 2f, brown) (Video 8)** while still being able to overcome obstacles. These results demonstrate the importance of rugged terrains in studying locomotor control: they expose the failure modes of controllers that otherwise work on flat terrain. Hereon, for our more complex sensorimotor tasks, we use the hybrid controller driven by a two-dimensional descending signal to control walking speed and turning by modulating oscillator frequencies and amplitude asymmetries, respectively **(Figure 2g)**.

### Vision and olfaction enable sensory navigation

To reach attractive objects (e.g., potential mates, food sources), avert from repulsive features (e.g., pheromones from predators), and avoid obstacles, animals use hierarchical controllers: higher-order brain systems must process sensory signals, use them to select the next course of action, and then transmit directives via descending pathways to lower-level motor systems. To simulate this sensorimotor hierarchy, we next added vision and olfaction to NeuroMechFly **(Figure 3a)**.

**Figure 3:**
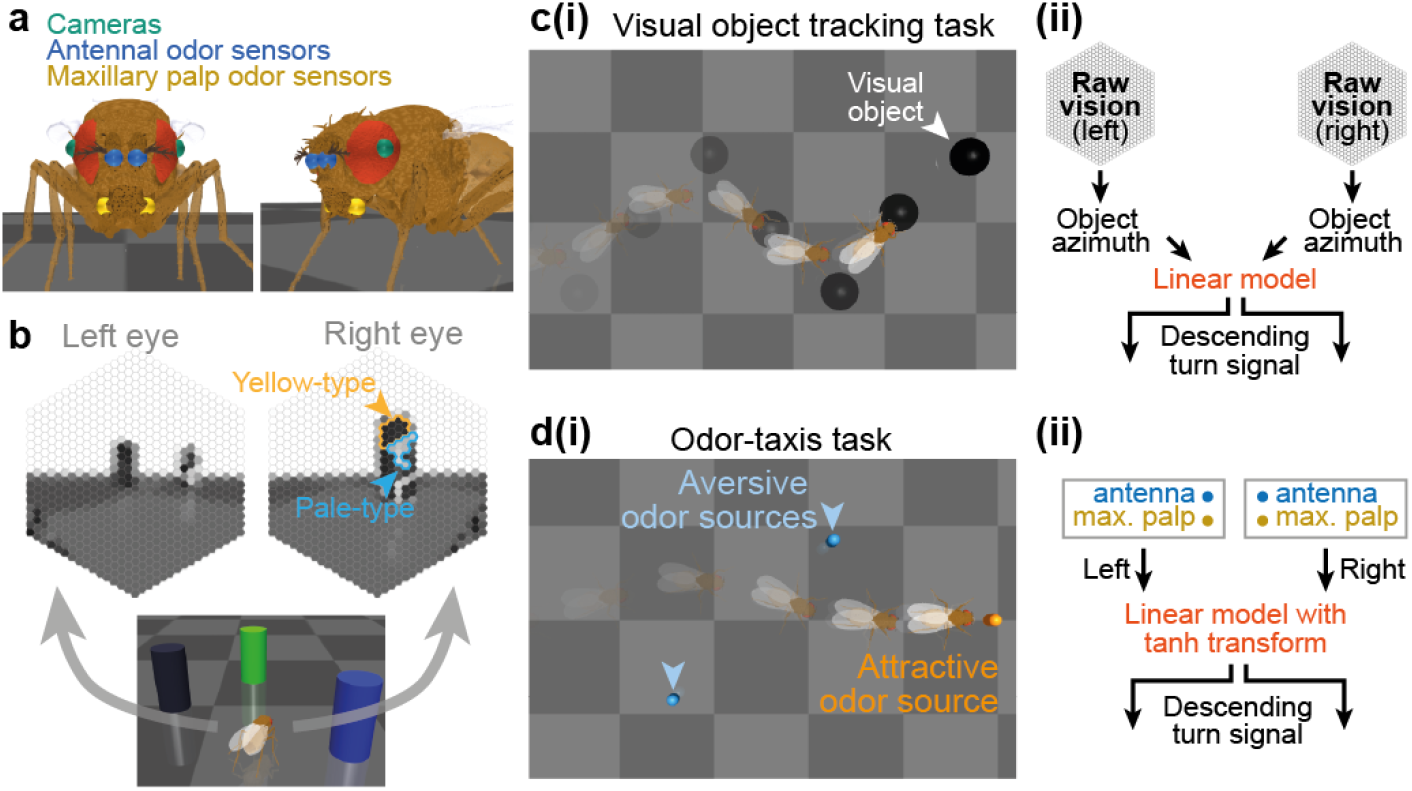
Vision and olfaction enable closed-loop sensorimotor control. **(a)** The placement of cameras for vision (green), and odor sensors—antennae (blue) and maxillary palps (yellow)—on the fly model’s head. **(b)** A simulation of the visual inputs perceived by the fly while facing three different colored pillars. Ommatidia perceive the same natural color in two different intensities due to selective sensitivity to different wavelengths of the two ommatidia types. For example, the blue pillar is perceived as darker by the yellow-type and as brighter by pale-type ommatidia. Indicated are some example yellow- and pale-type ommatidia. The positions of ommatidia types are randomly assigned. **(c)** A visual object tracking task. (left) The fly must follow a black sphere moving in an S-shaped trajectory. (right) To execute this task, we first extract the azimuth of the centroid of the object on the retina for each eye. These azimuths are then linearly transformed and capped to generate the appropriate descending command to CPG-based controllers. **(d)** An odor-taxis task. (left) The fly must seek an attractive odor source while avoiding two aversive odor sources. Note that the blue and orange odor markers are for visualization purposes only and are not seen by the simulated fly. (right) To execute this task, we first multiply the differences in mean odor intensities sensed by the antennae and maxillary palps on either side of the head by the gains of the corresponding odor types. The product is then passed through a nonlinear function restraining its range which is supplied to the descending controller.

A fly’s compound eye consists of ∼700–750 individual units called ommatidia that are arranged in a hexagonal pattern [43]. We emulated this by attaching a color camera to each of our model’s compound eyes **(Figure 3a, green)** and then transformed each camera image into 721 bins on a hexagonal grid [31] **(Figure 3b)**. We assumed a 270° combined azimuth for the fly’s field of view, with a ∼ 17° binocular overlap **(Extended Data Figure 3)**. As an initial step toward enabling heterogeneous color sensitivity in our model, we implemented yellow- and pale-type ommatidia—sensitive to the green and blue channels of images rendered by the physics simulator. Users can substitute the green and blue channel values with the desired light intensities sensed by yellow- and pale-type ommatidia to achieve more bio-realistic chromatic vision.

In addition to vision, we also made it possible for our model to detect odors in the simulation environment. Flies have olfactory receptor neurons (ORNs) in their antennae and maxillary palps. ORNs detect specific odorant molecules and convey this information to the brain’s antennal lobe, where it is further processed [44]. We emulated olfaction by attaching virtual odor sensors to our model’s antennae and maxillary palps **(Figure 3a)**. These virtual sensors can detect odor intensities across a multi-dimensional space that can be thought as representing, for example, the concentrations of monomolecular chemicals sensed by ORNs, or the intensities of composite odors co-activating numerous projection neurons in the antennal lobe. The modularity of our framework makes it possible for users to add more sensors to specific head locations and to implement additional signal processing by downstream olfactory centers (e.g., lateral horn or mushroom body [45]).

To illustrate the use of visual and olfactory sensing, we implemented visual object tracking and olfactory chemotaxis. In our object tracking task, the fly model had to visually track and follow a black sphere moving along an S-shaped trajectory in the environment. The controller processed the object’s visual location by computing its centroid position on the retina. Then, these visual features were linearly transformed into a two-dimensional descending signal **(Figure 3c, left)** that modulated the frequencies and amplitudes of CPG-based oscillators on each side of the body **(Figure 2g, “Turning controller”)**. This strategy allowed the fly to effectively track the sphere **(Figure 3c, right; Video 9)**. In our odor-seeking task, the simulated fly had to reach an attractive odor source while avoiding two aversive odor sources. The controller used sensors on the antennae and maxillary palps to compare the relative intensities of attractive and aversive odors across the left and right sides of the head [46]. These intensity values were multiplied by weights of opposite signs for attractive versus aversive odors. This left-right bias was used to asymmetrically control the descending signal **(Figure 3d, left)**, yielding effective odor-based navigation through the environment **(Figure 3d, right; Video 10)**.

### Ascending signals for path integration and head stabilization

Thus far we have demonstrated how brain-level sensory processing can drive the motor system via descending control. The inverse, ascending signals are thought to convey information back to the brain for action selection, motor planning, and sensory contextualization [47]. We next investigated how ascending feedback enables the modeling of important behaviors like path integration and head stabilization.

To effectively navigate the world, many animals, including flies [48], perform path integration wherein they constantly estimate and keep track of their own heading and distance traveled (“odometry”). The source of these idiothetic cues for path integration is unknown but may, in principle, be derived from ascending leg proprioceptive and tactile signals. We next explored how ascending proprioceptive and tactile feedback might be used to inform the brain of the change in body orientation and displacement. For each leg, we accumulated stride lengths by computing the forward translation of the leg tip relative to the thorax when the leg was in contact with the ground. Then, we computed the differences and sums of the left and right total stride lengths for each pair of legs within a short time window. These left-right differences and sums were used to predict the change in heading and forward displacement, respectively **(Figure 4a)**. Despite being linear, our model could give accurate predictions of the change in heading and forward displacement **(Figure 4b)**. These signals could be integrated over time to accurately estimate the fly’s true 2D position **(Figure 4c)**. We note, however, that the heading of the fly could sometimes be wrongly estimated **(Extended Data Figure 4)** in rare instances when the heading change prediction was off by a large margin. Thus, position estimates based on idiothetic cues alone can be prone to errors in heading integration even with an exceptionally well-performing internal model (*r*^2^ = 0.97, **Figure 4b**). This suggests that calibration using external sensory (e.g., visual) cues may be crucial.

**Figure 4:**
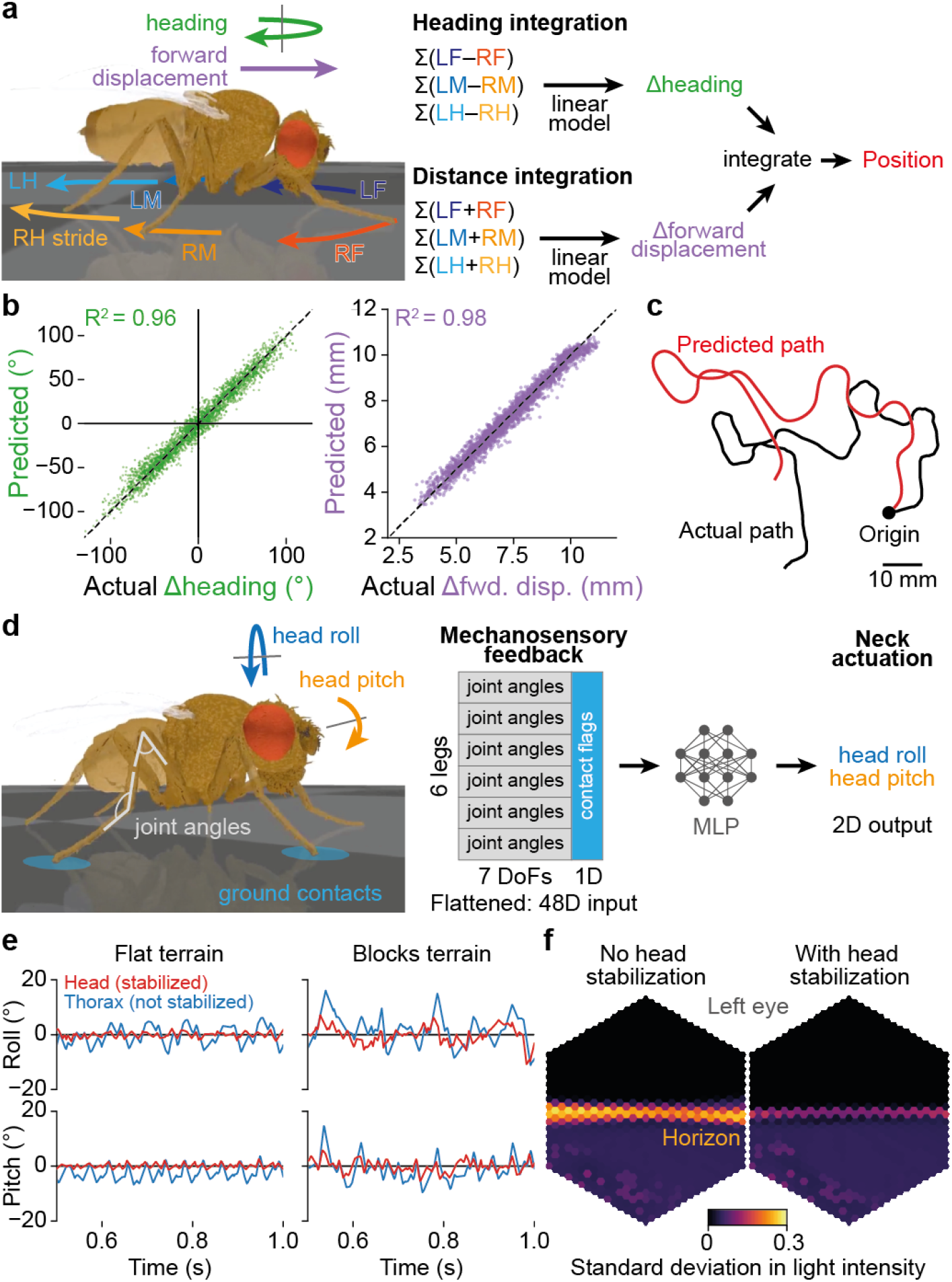
Ascending motor feedback for path integration and head stabilization. **(a)** Schematic of the path integration task. The simulated fly walks in a random trajectory over flat terrain. It must use proprioceptive signals from leg strides to predict the change in heading and forward displacement and integrate these signals over time to estimate its own position. **(b)** Scatter plot for 15 test trials (only 0.1% of all points are shown to facilitate visualization). Shown are the actual versus predicted changes in heading (left) or forward displacement (right). **(c)** An example of an actual versus predicted walking path. **(d)** Schematic of the head stabilization task. The fly model walks over flat terrain. A multi-layer perceptron (MLP) is trained to use leg joint angles and ground contacts to predict a neck roll and pitch actuation to compensate for thoracic movements and to stabilize the visual scene. **(e)** Time-series of head and thoracic roll and pitch with respect to the horizon when walking over either flat or blocks terrain. Shown are data with head stabilization. Note that without head stabilization the head and the thorax are coupled. **(f)** The standard deviation of ommatidia readings from the left eye while walking over flat, featureless terrain without (left) or with (right) head stabilization. Note the high variability in light intensity near the horizon when the head is not stabilized. This is due to more pronounced self-motion of the head.

In addition to informing path integration, ascending signals are well poised to perform other important tasks including visually tracking landmarks or targets (e.g., potential mates) while navigating over rugged terrain. In this context head stabilization may be controlled using leg sensory feedback signals [49] to compensate for body pitch and roll [50]. To explore this possibility in our embodied model, we designed a controller in which leg joint angles (i.e., proprioceptive signals) and ground contacts (i.e., tactile signals) were fed into a multi-layer perceptron (MLP). This MLP was trained to predict the appropriate neck joint actuation (i.e., head roll and pitch) required to cancel visual rotations caused by the animal’s own body movements during locomotion (using the hybrid controller) over either flat or blocks terrain **(Figure 4d)**. These predictions made from mechanosensory signals could indeed be used to dampen head movements in the roll and pitch axes compared to thoracic movements **(Figure 4e)** in a manner reminiscent of data from blowflies [50]. As a result, visual inputs, especially near the horizon, were more stable (i.e., exhibiting less variance in light intensity when walking in a featureless environment, **Figure 4f; Extended Data Figure 5a–b; Video 11**). Restricting the number and kinds of leg proprioceptive inputs to the MLP shows that ascending feedback concerning multiple degrees of freedom appears to be necessary to estimate body orientation for head stabilization **(Extended Data Figure 5c–d)**.

### Hierarchical controller trained with reinforcement learning

With the elements of a hierarchical controller in place, it becomes possible to leverage modern machine learning approaches to train a network to accomplish more complex tasks. To illustrate this, we trained our fly model to avoid an obstacle while searching for an attractive odor source over rugged terrain. In total, our hierarchical controller [16] **(Figure 5a)** consisted of: (i) a vision module (a convolutional neural network on a hexagonal lattice) that extracts the object’s direction, distance, locations, and sizes on the retinas **(Extended Data Figure 6)**; (ii) a decision module (an MLP) that receives as inputs preprocessed visual features and odor intensities from each antenna and computes a turning bias; (iii) a twodimensional descending signal that modulates locomotor CPGs that drive walking and turning; (iv) a hybrid walking controller **(Figure 2g)**; and (v) an ascending feedback module that performs head stabilization. We trained the vision module in a supervised manner by randomly placing the fly and obstacle in the arena to collect training data and trained the decision module through reinforcement learning. This hierarchical controller could achieve multi-modal visual-olfactory navigation over rugged terrains **(Figure 5b-c; Video 12)**. This integrative task demonstrates how one can define individual components in a modular fashion and combine them to investigate a hierarchical sensorimotor task in closed loop using NeuroMechFly v2.

**Figure 5:**
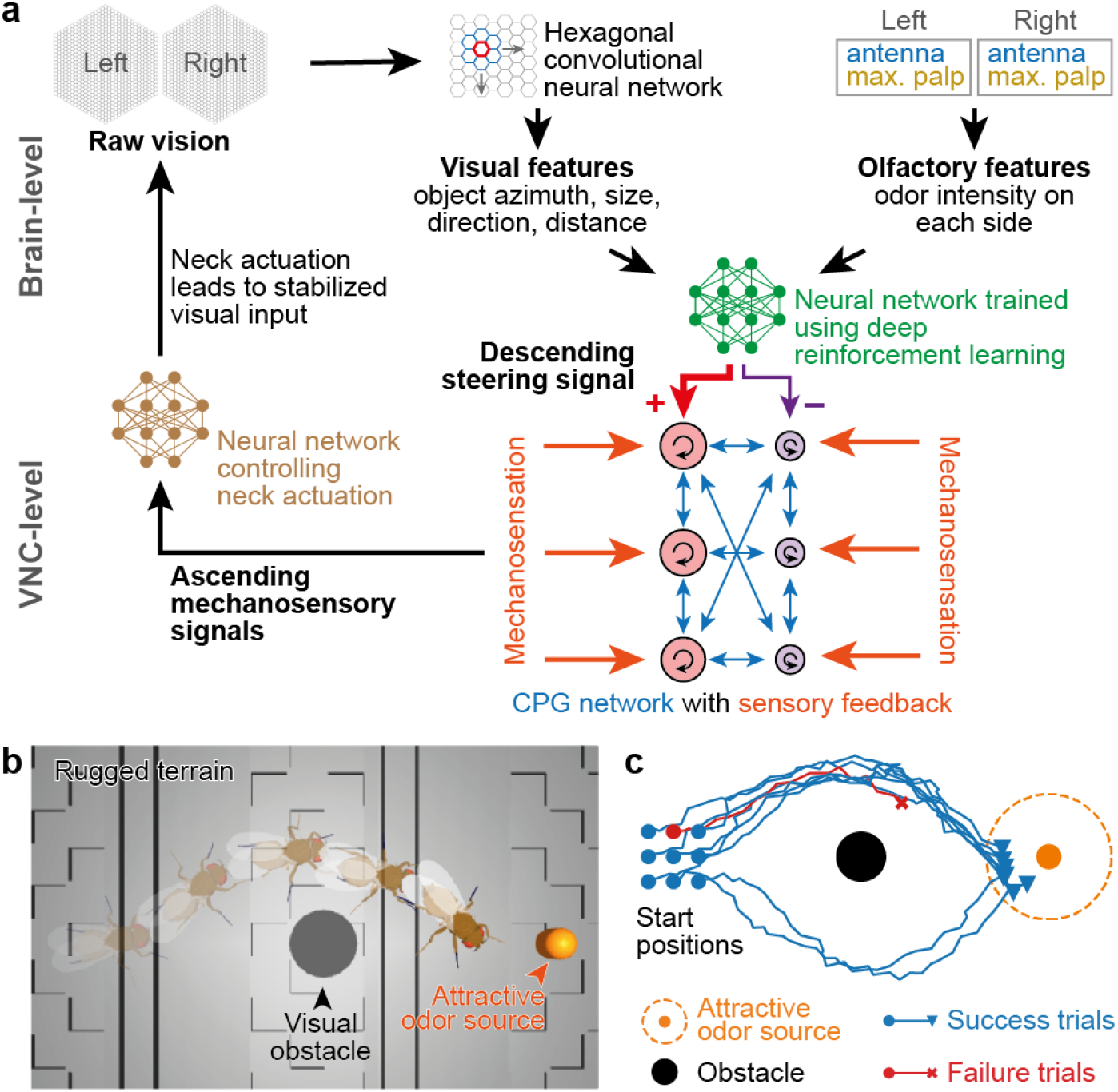
Using reinforcement learning to train a hierarchical controller for a multi-modal task. **(a)** Schematic of the multi-modal reinforcement learning-based navigation controller. Visual features are extracted using a convolutional neural network. Olfactory features are processed as in **Figure 3d**. These features are input to an artificial neural network which is trained through reinforcement learning to output appropriate descending turning commands. These commands are executed by a hybrid low-level motor controller integrating CPGs and sensory feedback and using leg adhesion. Ascending motor feedback signals are used to perform head stabilization for visual (and to a lesser extent olfactory) processing. **(b)** The decision-making network is trained using reinforcement learning to enable the fly to seek an attractive odor source while avoiding a visually detected obstacle over complex terrain. An example of one successful trial is shown. The orange odor marker is for visualization purposes only and is not visible to the simulated fly. **(c)** The trajectories of the fly in 9 examples (8 successful; 1 failed) beginning from different spawn positions (circle markers).

### Using more bio-realistic algorithms for sensorimotor control

With full access to raw light intensities and odor concentrations, users can build their own sensory-rich environments and process these with models of even higher levels of biological realism. We first illustrate this by using a *Drosophila* olfactory taxis algorithm to navigate a complex odor plume. We simulated a plume embedded in airflow **(Figure 6a)** representing the propagation of, for example, an attractive food odor in the real world. Unlike the simplified controller used in **Figure 3d**, here we implemented a previously proposed *Drosophila* plume navigation algorithm [51] to control locomotion. Using this algorithm, to reach the odor source, the fly randomly switches between forward walking, pausing, and turning **(Figure 6b)**. These actions are governed by Poisson processes, with Poisson rates and turning direction bias modulated based on odor encounters to favor navigation towards the target. In our simulation, the fly successfully reached within 15 mm of the odor source **(Figure 6c, Video 13)** with a low (9 out of 100 trials), albeit similar success rate comparable to what was seen in real flies in a larger arena over a longer time period [51].

**Figure 6:**
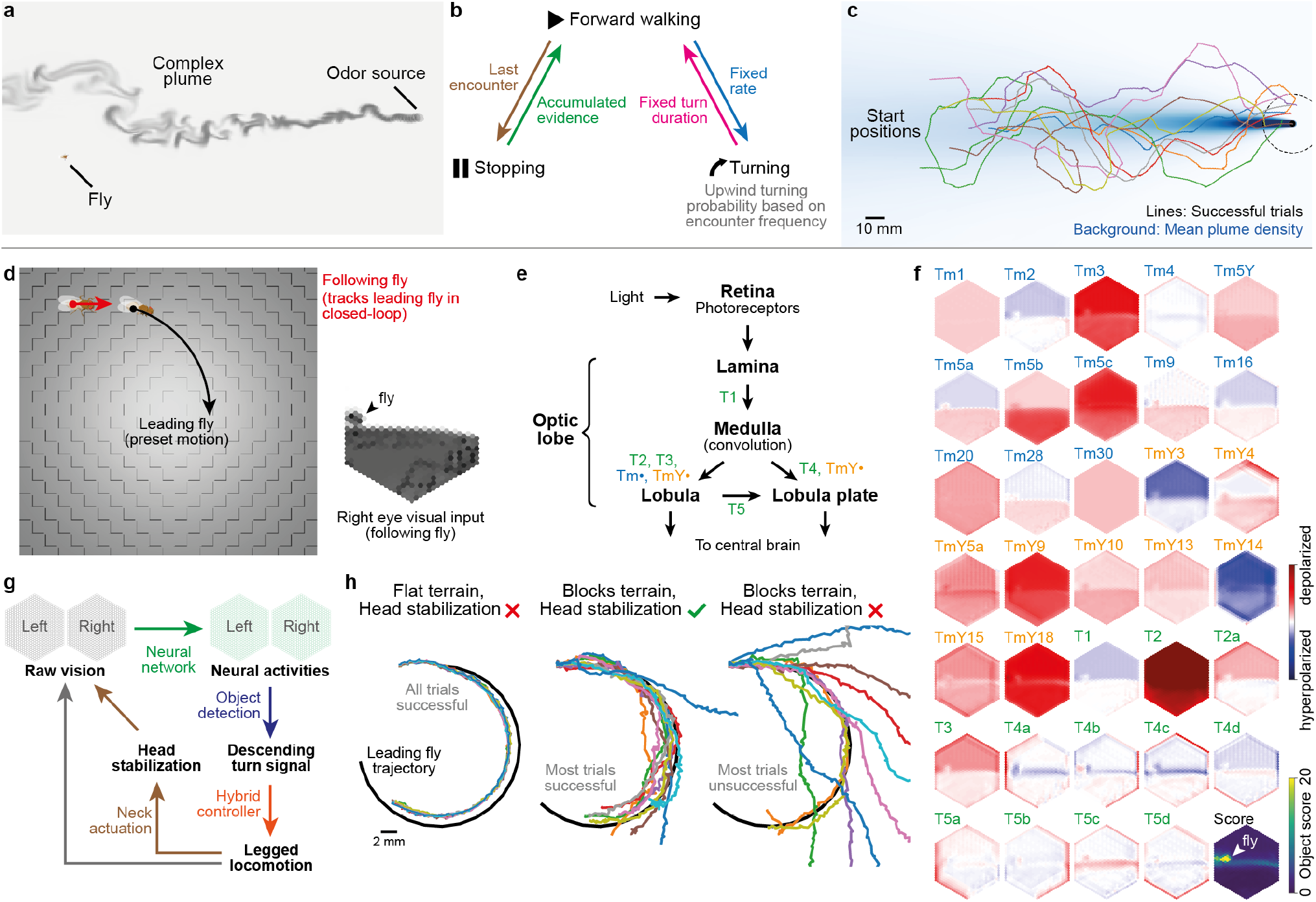
Incorporating more bio-realistic environments and controllers. **(a)** Modeling a complex odor plume to simulate taxis toward an attractive odor source (from [51]). **(b)** Transitions between locomotion states based on plume encounters. The transition from forward walking to stopping is governed by a Poisson process whose rate changes based on the time since the last encounter. The transition from stopping to forward walking is governed by a Poisson process whose rate changes based on a history of recent encounters (accumulated evidence). The transition from forward walking to turning is governed by a Poisson process with a fixed rate. Each turn has a fixed duration. The probability of turning upwind increases as the encounter frequency increases. **(c)** Fly trajectories for 9 successful trials when the odor source (dashed circle) was reached. Trajectories are color-coded. The mean plume density is shown in blue on the background. **(d)** Overview of the fly following task. Two fly models are spawned. The first (“leading”) fly has a preset circular walking trajectory. The second (“following”) fly uses a hierarchical controller to follow the leading fly. **(e)** Block diagram of the *Drosophila* optic lobe. Indicated in blue and green are the neurons used to detect the leading fly. Only connections directly relevant to the neurons used in the fly-tracking task are shown. **(f)** Object detection score and activity patterns of 34 putative output columnar neuron types (of 65 total) drawn from a connectome-constrained visual system network [31] when the *leading* fly is moving in the field-of-view of the *following* fly’s right eye. **(g)** Schematic of a hierarchical controller using visual inputs, a connectome-constrained visual system network, processing of population activity, object detection, descending control of a hybrid controller with leg adhesion, and ascending feedback for head stabilization. **(h)** Performance of this more bio-realistic visual system network on fly following with or without ascending feedback-based head stabilization. Shown are 11 trials each.

Ultimately, to gain insights into how the real fly brain works, one would explore controllers with artificial neurons that can be mapped to real neurons or neuronal cell types. This may be achieved by building artificial neural networks with architectures constrained by the connectivity of the brain [24] and the VNC [25, 26]. To illustrate how such models might be embodied and studied in the context of autonomous behavior, we designed a “fly following” task in which a fly must use a realistic visual system model **(Figure 6d)** to follow another fly—akin to chasing behaviors during courtship. We used a recently constructed connectome-constrained model [31] for this task. We interfaced NeuroMechFly with this visual system model to emulate layered visual processing in the fly brain. Concretely, we passed the visual experience of the *following* fly as inputs to this pretrained connectome-constrained model and used the activities of T1–T5, all Tm, and all TmY neurons (indicated as putative output cells types [31]) to perform object detection **(Figure 6f)**. Then, based on the position of the detected object, we modulated the descending turning signal **(Figure 6g)** to drive the hybrid controller **(Figure 2g)**, which controls walking. We asked to what extent ascending feedback-driven head stabilization **(Figure 4d)** is necessary to enable reliable fly following. We found that, although over flat terrain the *following* fly could successfully track the *leading* fly without head stabilization **(Extended Data Figure 7)**, stabilization was crucial for tracking over rugged terrain **(Figure 6h, Video 14)**. We obtained similar results even when using a smaller subset of neurons that provide inputs to LC9 and LC10–LC neurons implicated in courtship in particular [52] **(Extended Data Figure 7)**. These results highlight how realistic neural networks can be coupled with embodied models to close the sensorimotor control loop.

## Discussion

Here we have introduced NeuroMechFly v2, a framework for performing integrated sensorimotor neuromechanical simulations of the adult fly, *Drosophila melanogaster*. In **Supplementary Note 1**, we provide a summary of general features currently supported in NeuroMechFly v2, specific modeling choices concretely demonstrated in this paper, and opportunities for future work. Because our simulation framework is modular, researchers can build integrated models in an interoperable manner by choosing the appropriate level of detail for each part of the model to suit the scientific question under consideration. For example, one can use more abstract baselines or existing models for control elements outside the focus of investigation. Although important behaviors like those involving the control of wings/halteres (e.g., flight [14]), abdomen (e.g., egg-laying), and proboscis (e.g., feeding) are not yet implemented, they are supported within this framework. With enriched sensory feedback and improved biomechanics, NeuroMechFly v2 enables the whole-body simulation of complex behaviors requiring controllers that span sensing, navigation, internal states [22], learning [45], and motor control.

In the future, our simulation framework is likely to be further improved in a number of ways. First, we anticipate that recent developments in physics simulation, particularly GPU acceleration and differentiable simulation will facilitate the training of larger models through reinforcement learning. Second, careful measurements and analyses of the *Drosophila* musculoskeletal system (i.e., tendons and muscles) could improve the interface between neural network controllers and the biomechanical embodiment. Third, as additional connectome-constrained neural circuit models become available, they can be added to the corpus of controllers in our modular simulation framework. FlyGym’s compatibility with the Gymnasium API will ensure that changes are implemented relatively easily without disrupting the established user interface. In the more distant future—following significant improvements in modeling infrastructure enabling high-throughput, low-latency simulations—a similar simulation framework could be integrated into closed-loop experiments. For example, NeuroMechFly could be used during experiments to replay an animal’s kinematics as captured by pose estimation methods, enabling the real-time inference of dynamic variables such as contacts and informing experimental perturbations in closed loop [53]. These efforts will bring the field closer to achieving the ultimate goal of uncovering neuromechanical mechanisms giving rise to adaptive animal behaviors in a sensory-rich and physically complex world.

## Acknowledgments

We thank Victor Lobato-Rios for valuable insights and early exploration of visual inputs to the model. We thank Jonathan Arreguit, Shravan Tata Ramalingasetty, and Auke Jan Ijspeert for the development of FARMS, which was used to generate the MJCF file of the updated model. We thank Janne K. Lappalainen, Jakob H. Macke, Srinivas C. Turaga and colleagues for making the FlyVision model available before publication. PR acknowledges support from a Swiss National Science Foundation (SNSF) Project Grant (175667) and an SNSF Eccellenza Grant (181239). SWC acknowledges support from a Boehringer Ingelheim Fonds PhD fellowship. PGÖ acknowledges support from a Swiss Government Excellence PhD Scholarship and a Google PhD Fellowship. FH acknowledges support from a Boehringer Ingelheim Fonds PhD fellowship.

## Author contributions

S.W.C. — Conceptualization, Methodology, Software, Formal Analysis, Investigation, Data Curation, Validation, Writing – Original Draft Preparation, Writing – Review & Editing, Visualization.

V.A.S. — Conceptualization, Methodology, Software, Formal Analysis, Investigation, Data Curation, Validation, Writing – Original Draft Preparation, Writing – Review & Editing, Visualization.

T.K.C.L. — Conceptualization, Methodology, Software, Formal Analysis, Investigation, Data Curation, Validation, Writing – Review & Editing, Visualization.

P.G.Ö . — Conceptualization, Methodology, Software, Investigation, Data Curation, Validation, Writing – Original Draft preparation, Visualization, Writing – Review & Editing.

L.G. — Methodology, Software, Validation, Investigation, Writing – Review & Editing.

F.H. — Conceptualization, Methodology, Software, Investigation, Writing – Review & Editing.

P.R. — Conceptualization, Methodology, Resources, Writing – Original Draft Preparation, Writing - Review & Editing, Supervision, Project Administration, Funding Acquisition.

## Methods

### The FlyGym package

FlyGym is implemented based on MuJoCo [33] and the dm control [54] library and complies with the Gymnasium API [5] for the Markov Decision Process. The user interacts with the simulation through *actions* and *observations* **(Figure 1, Supplementary Note 2)**. The definition of the action and observations in the default control task can be found in **Supplementary Note 3**. More detailed, low-level information can be accessed directly using dm control and MuJoCo.

We configured the meshes at 1000 *×*scale in MuJoCo to obtain observation measurements in mm and mN. The user can implement preprogrammed premotor computations and sensory preprocessing by extending the base `Fly or Simulation` classes **(Figure 1c)**. This will modify the action and observation spaces accordingly.

If the user wishes to use a simplified, ball-and-stick model to speed up computation, one can use a MuJoCo feature that approximates body geometries as “capsules” (i.e., cylinders with a hemisphere at each end). To do so, the user should replace `type=“mesh” with type=“capsule” in the <geom>` tag of the model MJCF file.

### Updated rigging of the biomechanical model

In *Drosophila*, the antenna consists of three main segments—the scape, pedicel, and funiculus—in addition to the arista [55]. The fly has four muscles that can actively control the joint between the scape and pedicel [56]. By contrast, the funiculus and the arista move or deform passively in response to external forces (e.g., wind, limb contact during grooming). In the original NeuroMechFly model, the entire antenna could move relative to the head with one DoF. We improved the model by separating each antennal mesh into three different meshes using Blender. In the biomechanical model, “bones” determine how objects move with respect to one another. We positioned the bones to accurately replace joints based on anatomical features such as the stalk-like structures connecting the funiculus to the pedicel [57]. We then constructed a kinematic chain connecting these segments: scape-pedicel-funiculus-arista from proximal to distal. Instead of simulating the arista as a soft body (which is computationally expensive), we emulated the compliance of the arista by adding three DoFs between the funiculus and the arista. The passive movement of the arista can be fine-tuned by modifying the stiffness and damping coefficients of these DoFs. We gave the remaining joints (i.e., head-pedicel and pedicel-funiculus) all three rotational DoFs because the real number of DoFs in these antennal joints remains unknown. Future users can modify each DoF (e.g., fix/unfix or stiffen) in the model file to emulate the measured dynamics of the antennae.

The position of the neck joint affects the translation of segments on the head, such as the proboscis, antennae, and eyes. The neck is located ventral to the hair plate behind the head. In our previous model, the neck had one unactuated pitch DoF. Here we modified the location of the neck joint by comparing the head rotations of the model with those of the real fly and added two known DoFs (yaw and roll) to the neck. Furthermore, we spaced the head away from the thorax to emulate the space filled by the neck. The size of the neck was determined by measuring the proportion of head size to neck size in real animals [58]. We confirmed that the rotation center of the neck joint fits the original pose from the NeuroMechFly CT scan by actuating the neck joint to match the original pose. Next, we adjusted the positions of the front legs based on the distance between the front leg thorax-coxa position and anatomical landmarks (e.g., humeral bristles) and an overlay of camera images of real animals with images of the model. Finally, we changed the resting pose of the model such that the angle of the scutellum would resemble that of real animals standing freely (untethered) on flat terrain. We used the FARMS simulation framework [59] to generate the MJCF file of the updated model.

### Leg adhesion and critical climbing angle

Leg adhesion was added using built-in MuJoCo actuators. Adhesion takes the form of an artificial force injected normal to the point of contact. This force is oriented toward the object colliding with the body part containing the actuator. If multiple contacts occur with external objects and the adhesion actuated body, the force is equally divided between these contact points.

In our model, adhesion is actuated and can be turned on and off during locomotion. We manually defined the adhesion on/off periods within the preprogrammed stepping pattern **(Extended Data Figure 2)**. Adhesion is on during the stance phase and off during the swing phase **(see “Stepping pattern”)**. We controlled adhesion in a binary fashion but it is possible to use a gradient of adhesion forces by modulating the input to the adhesion actuator at every time step.

To quantify the impact of maximal adhesive force on the ability of the fly model to climb **Figure 2c**, we measured the critical slope—the angle in degrees at which the fly could no longer maintain forward locomotion, or flipped—as a function of the maximal adhesion force. Flipping is defined as when either the absolute roll or pitch angle of the fly is above *π*/2. A fly has failed to maintain forward locomotion if its position along the surface is negative compared to its initial position after 1 s.

### Stepping pattern

We derived the kinematics for each individual step from manually annotated video recordings of a real fly during untethered walking. We recorded bouts of straight walking at 360 Hz in a linear chamber (12 mm long *×* 4 mm wide *×* 2 mm tall) with prisms as walls. The video was down-sampled to 120 Hz. Then five leg keypoints (thorax-coxa, coxa-trochanter, femur-tibia, tibia-tarsus joints, and claw), both antennae, neck, thorax, and abdomen keypoints were manually annotated from a 0.3 s episode of straight walking. The recording was performed on a wildtype (PR) female adult *Drosophila melanogaster* raised at 25 °C and 50% humidity on a 12-hour light-dark cycle. The fly was recorded 4–5 days post eclosion.

We determined the 3D position of each keypoint assuming that the prisms are oriented at 90°. We aligned resulting 3D poses to the template of NeuroMechFly’s skeleton by scaling the length of the full leg. Finally, we applied inverse kinematics to each kinematic chain to obtain joint angles [60]. From the recording we then segmented eight swing-to-swing steps and eight stance-to-stance steps. Out of these 16 unique steps (five front leg steps, six middle leg steps, and five hind leg steps) seven were insufficiently closed (i.e., mean distance between the first and last joint angle in the step greater than 0.17, 0.12, and 0.17 radians for the front, middle and hind legs, respectively) and were therefore discarded. The final stepping pattern is composed of three steps that are of the same lengths, are closed, and, when mirrored, yield symmetric and smooth steps.

We obtained the final stepping pattern by (i) segmenting each of the nine selected steps, (ii) stretching or compressing them to the median step length of 0.135 s, (iii) linearly interpolating the difference between the first and last time-points of the step through the last 10% of each step to guarantee perfect closure, (iv) modifying all steps so that phase 0 corresponded to the initiation of the swing, and (v) finally generating a complementary dataset with mirrored joint angles so that each step could be replayed in either right or left legs irrespective of their leg of origin. From those nine steps, we obtained 30 combined stepping patterns. We used the combination that maximized the displacement along the fly’s initial heading direction and minimized its lateral displacement.

### CPG-based controller

Central Pattern Generators (CPGs) are neural circuits that generate rhythmic outputs without receiving rhythmic input [40]. Through interactions, coupled CPGs can synchronize with given phase offsets. As for the previous version of NeuroMechFly, we implemented CPGs by adapting those used to model limb actuation in salamanders [4]. All DoFs for a given leg of the fly were controlled by a single CPG. The oscillatory output of a given CPG was then interpreted as the phase and amplitude of the step cycle. The gait pattern emerges from the phase biases of the different CPGs. We used an idealized tripod gait for walking in our CPG model. The precise definition of the CPG network can be found in **Supplementary Note 4** and the parameters are detailed in **Supplementary Tables 2–3**.

### Rule-based controller

We used a rule-based controller to illustrate a decentralized control architecture. This controller was inspired by the first three rules described in Walknet [1, 61, 62]. The first rule ensures stability by inhibiting swing onset in the rostral neighbor of a swinging leg. The second rule ensures the propagation of the wave by eliciting a swing in the rostral and contralateral neighbors of a leg entering stance phase. The third rule enforces temporal coherence by eliciting a swing in the caudal and contralateral neighbors of a leg approaching the end of its stance phase. The rules modify a stepping likelihood score for each leg, and a step is initiated on the leg in stance phase with the highest positive score. If all legs have negative scores, no step is initiated. If multiple legs have similar scores (difference *<* 0.1% of the highest score), a leg is selected at random to avoid artifacts resulting from small numerical errors. The contributions of these rules are weighted **(Supplementary Table 4)**: rule 1 is weighted most heavily as it is crucial to maintain stability. Rules 2 and 3 are given different weights for ipsi- and contralateral connections. To maintain synchrony, we ensured that the duration of the swing and stance periods were identical across all legs. To more fairly compare the rule-based controller with the CPG controller, we scaled the duration of steps to match the stepping frequency of the CPG controller.

### Hybrid controller

The hybrid controller is a CPG controller with two additional rules that can be activated depending on leg mechanosensory signals. These rules allow the fly to recover when a leg becomes stuck in a gap (e.g., in gapped terrain) or hits an obstacle (e.g., in blocks terrain) by adjusting the leg in question. The first rule (“overstretch rule”) is activated when a leg is extended farther than expected along the z-axis (indicating that the leg may have fallen into a gap). More precisely, this rule becomes active when the tip of a leg is *>*0.05 mm lower than the third most extended leg along the z-axis. Due to numerical errors and physics instabilities, the z-positions of the tips of the legs read out from the physics simulator are sometimes slightly below 0 when the legs are on the ground. A 0.05 mm margin was therefore added to avoid spurious detection of leg overstretch. If multiple legs meet this criterion, only the leg that extends the furthest is corrected. The second rule (“stumbling rule”) is activated when a leg comes into unexpected contact with an object, resulting in a horizontal force against the direction of locomotion. More precisely, this rule becomes active when the tibia or the two most proximal segments of the tarsus have a contact force greater than 1 mN opposing the heading of the fly while the leg is in swing. When either rule is activated, a shift is progressively added to a subset of joints on the leg in question such that the leg lifts up higher than normal during swing or extends slightly more during stance. The step phase dependence of the adjustment is obtained by using a gain described by a piece-wise linear function reaching a maximum of 0.8 at the swing midpoint, a minimum of -0.1 at the stance midpoint and remaining at 0 from the beginning of the swing to the end of the swing plus one-eighth of a cycle. **Supplementary Table 5** provides a summary of the joints involved in leg retraction and their rates of change. Both rules are persistent to ensure proper release of the leg: if one rule was active during the past 0.002 s, the leg enters a persistence period prolonging the adjustment for 0.002 s. Once the persistence period is over and as long as the rules are no longer active, joint angles progressively reset. To avoid over-correction, the leg’s position is adjusted for 0.008 s before the increment is capped. The swing duration was extended by one-eighth of a cycle to delay the initiation of adhesion and give more time for the leg to clear any obstacles.

### Benchmarking of locomotor controllers over rugged terrains

We benchmarked locomotor controllers by running 20 simulations, starting from different spawn positions and initial states, for 1.5 s each and computing the average velocity in the horizontal plane. Walking speeds driven by the controllers are comparable because the same preprogrammed step is used for all controllers and walking speed is only influenced by inter-leg coordination. “Gapped terrain” consists of horizontal 1 mm-wide blocks separated by 0.3 mm-wide, 2 mm-deep gaps. “Blocks terrain” consists of 1.3*×*1.3 mm blocks configured in a checkerboard pattern, with half of the blocks 0.35 mm higher than the others. A small overlap is added between blocks to avoid extremely thin surfaces near the corners that can lead to physics instabilities. “Mixed terrain” consists of alternating flat, gapped, and block patterns along the x-axis. We used a joint position gain *k*_*p*_ of 45 and an adhesion force of 40 mN for all controllers.

### Control of turning

Walking flies execute turns on a continuum of sharpness. The turning program is controlled by descending neurons [63]. For smoother turns (i.e., rotations *<* 20°) the fly mostly increases the stroke amplitude of its outer legs. For sharper turns (20°–50°) the fly additionally decreases the stroke amplitude of its inner legs. For very sharp in-place turns (*>* 50°), the fly steps its inner legs backward [64]. Our controller receives a two-dimensional descending input that controls turning **(Figure 2g)**. On each side, the descending signal DN ∈ ℝ modifies the intrinsic frequency *ν*_*i*_ and maximum amplitude *R*_*i*_ of each oscillator *i* as follows:

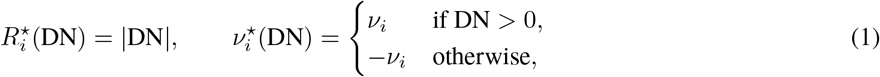

where 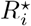 and 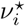 are the modified maximum amplitude and intrinsic frequency.

### Implementation of vision

Flies have three major types of ommatidia—units arranged in a hexagonal pattern to make up the compound eye. These differentiate colors and polarization properties by using different combinations of photoreceptors. Yellow- and pale-type ommatidia are stochastically arranged throughout the eye and enable two-dimensional chromatic sensitivity in the UV _∼300 nm_-to-yellow range [65]. The yellow- and pale-type ommatidia are found at a 7:3 ratio [66]. A third type is found in the eye’s dorsal rim area (DRA) facing the sky and is specialized for polarization detection during navigation [67]; this type of ommatidia is not implemented in our model. The field of view of each eye is defined based on prior studies [68, 69]. In our implementation, yellow- and pale-type ommatidia are instead made sensitive to the green and blue channels of the physics simulator. For a more biologically accurate representation of color, the green- and blue-channel display colors can be set as the inner products of the actual surface reflectance spectrum of the object and the spectral response curves of the appropriate photoreceptors [65]. We corrected the input images to superimpose a “fish-eye” effect that makes the representation of angles consistent throughout the field of view **(Supplementary Note 5, Extended Data Figure 3c–d)**.

### Visual object tracking task

In our visual object tracking task, the fly follows a black sphere moving in an S-shaped trajectory at 10 mm/s. To achieve this, we first used a thresholding rule to detect the object (normalized light intensity 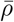 below 0.2). Then, we computed the position and size of the object (both normalized) as seen from each eye. Finally, we linearly adjusted the descending signal on each side depending on the object’s azimuth as seen from the ipsilateral eye. The turning bias is updated every 0.05 s of simulated time. More precisely,

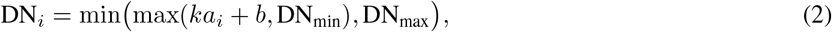

where DN_*i*_ is the descending signal on side *i*; *a*_*i*_ is the azimuth expressed as the deviation from the anterior edge of the eye’s field of view, normalized to [0, 1]; *k* = −3, *b* = 1 describe the response curve; DN_min_ = 0.4, DN_max_ = 1.2 are the minimal and maximal allowed values for the descending signal.

### Olfactory chemotaxis task

In the olfactory chemotaxis task, the fly seeks an attractive odor source while avoiding two aversive odor sources. To achieve this, we first calculate the odor intensities sensed at the locations of the antennae and maxillary palps based on a diffusion function, *I*(*d*), where *d* is the distance from the odor source and *I*(*d*) gives the odor intensity. The odor diffusion relationship can be defined by the user. In this example, we used the inverse square relationship *I*(*d*) = *I*_peak_/*d*^2^ where *I*_peak_ is the peak intensity. If there are multiple sources for the same odor, their intensities are summed. Then, for the attractive odor, we averaged intensities sensed by the antennae and the maxillary palps weighted by 9:1 (roughly comparable to the ratio of ORNs in the antennae and maxillary palps [70]). By contrast, to demonstrate the possibility of using different sensors for different odors, we used only the intensity sensed by the antennae for the aversive odor to emulate odorants that can only be sensed by one but not both organs. We performed this process for olfactory organs on each side of the head and multiplied the relative differences in intensities between both sides with a gain factor. Next, we summed up this product for each odor and nonlinearly transformed it into a turning bias. This bias modulates descending signals that drive turning. The turning bias is updated every 0.05 s in simulated time. More precisely,

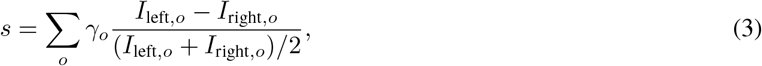

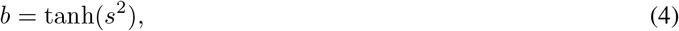

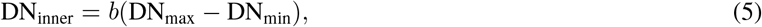

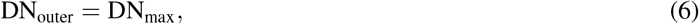

where *s* is the weighted sum of bilateral differences in odor intensities, *b* is the nonlinearly transformed turning bias, *I*_side,*o*_ is the mean intensity of odor *o* sensed by the antenna and the maxillary palp on the specified side; DN_inner_, DN_outer_ are the DN drives on the inner and outer sides (when *s >* 0, the fly performs a right turn, vice versa); DN_min_ = 0.2, DN_max_ = 1 define the range of the DN drives; and *γ*_*o*_ is the gain of odor *o* (*γ*_attractive_ = −500 and *γ*_aversive_ = 80).

### Path integration using ascending feedback

To test the degree to which path integration can be performed using ascending feedback, we constructed a scenario in which the fly model performs random exploration of a featureless environment and tries to estimate its position. To collect training data, we make the fly alternate between forward walking and in-place turning. Turning is modeled as a Poisson process with a rate *λ*_turn_ = 2 s^−1^. We deliberately chose a relatively high *λ*_turn_, compared to the range of typical fly behavior, to make path integration more difficult. We simulated walking using three walking gaits: tripod gait (3 legs in stance at a time), tetrapod gait (4 legs in stance at a time), and wave gait (5 legs in stance at a time). When the fly executes a turn, we apply a fixed asymmetrical descending drive of [DN_inner_, DN_outer_]. For the tripod and tetrapod gaits, [DN_inner_, DN_outer_] = [ *−*0.2, 1.0]; for the wave gait, [DN_inner_, DN_outer_] = [0.4, 1.0]. These choices led to qualitatively similar turning. The direction of the turn is chosen at random. The duration of the turn (and therefore the angle turned) is sampled from a normal distribution centered at 0.4 s with a standard deviation of 0.1 s. The fly receives no visual information—akin to navigating in the dark. We ran *N* = 15 trials with different random seeds for each of the three gaits. Each trial was 20 s long. For simplicity, the correction rules in the hybrid controller were disabled for this experiment. Then, we developed a path integration algorithm that separately predicts the changes in heading and forward displacement using the difference and sum of the cumulative stride lengths on the left and right sides. These signals are then integrated over time to estimate the fly’s position. Parameters in this algorithm are fitted to the aforementioned training data. A detailed description of the algorithm can be found in **Supplementary Note 6**.

### Head stabilization using ascending feedback

We first simulated walking over the “blocks” terrain and recorded movements of the thorax. We recorded joint angles and ground contacts throughout the simulation and calculated the optimal neck roll and pitch angles that would “cancel out” thoracic rotations. We used these angles as ground truth. Using the joint angles and ground contacts as inputs, we trained an artificial neural network (multi-layer perceptron, MLP) to predict the optimal correction angles. These predicted correction angles are then used to actuate the neck joint using a proportional derivative (PD) controller. Details of this process can be found in **Supplementary Note 7**.

### Multi-modal navigation task solved using reinforcement learning

In the multi-modal navigation task, the fly locomotes over rugged terrain to seek an attractive odor source while avoiding a visual obstacle in its path. To achieve this, we used a hierarchical controller consisting of (i) a vision module that extracts lower-dimensional visual features from retinal inputs, (ii) a decision module that predicts the appropriate turning bias given pre-extracted visual features and odor intensities, and (iii) a descending interface passing the turning bias to a downstream (iv) hybrid motor controller that integrates CPG states with leg mechanosensory feedback. To reduce training time, we slightly simplified the “mixed terrain” by reducing the gap width to 0.2 mm and the block height to 0.3 mm.

We started by training a convolutional neural network to extract features from the raw visual input, namely the direction of the object relative to the fly, the distance of the object from the fly, whether the object is within the fly’s field of view, the azimuth of the object seen from each eye, and the size of the object on the retina. Details of this vision preprocessing model can be found in **Supplementary Note 8**. Then, we trained an MLP to perform the multi-modal navigation task using Soft Actor-Critic (SAC), a reinforcement learning algorithm [71]. Details of the reinforcement learning task are outlined in **Supplementary Note 9**.

### Tracking complex odor plumes

We simulated the complex odor plume using PhiFlow [72] based on existing open-source software [73]. Once simulated, we provided the plume concentration *s* at the appropriate location to the odor sensors of the simulated fly. The plume is also overlaid onto the rendered image when applicable. Then, we implemented a tracking algorithm similar to the one in [51] wherein the fly makes decisions based on discrete, binarized plume encounters. Briefly, the fly switches between walking and stopping governed by Poisson processes whose rates depend on the time since the last odor encounter and the accumulation of odor encounters, respectively. The fly also turns based on a Poisson process where the direction of turning depends on the encounter frequency. Details about the odor plume simulation and the odor tracking algorithm can be found in **Supplementary Note 10**.

### Closed-loop fly following using a connectome-constrained visual system model

To simulate the responses of visual system neurons to the visual experience of the simulated fly, we interfaced NeuroMechFly with a published connectome-constrained visual system model [31] implemented in the FlyVision package (https://github.com/TuragaLab/flyvis). We modified its implementation to handle one frame at a time rather than the whole dataset at once, enabling closed-loop deployment. Since both FlyGym and FlyVision use a hexagonally arranged ommatidia grid with a side length of 16 ommatidia, there is a one-to-one mapping for ommatidia between the two packages. We sampled the visual experience of the simulated fly at 500 Hz and simulated the FlyVision model at the same rate. The two eyes are simulated independently. We used the best-performing model reported in the visual system modeling study [31] rather than the whole ensemble of models.

From the visual system simulation, we read out the activities of the 34 T-shaped transmedullary neurons, which are considered the outputs of the optic lobe. We compared the activities of these neurons with their baseline activities, obtained by simulating the fly walking in an empty arena. We then detected the object by identifying ommatidia for which neural activities differed significantly from baseline. More precisely, for each cell in the hexagonal ommatidia grid, we computed an object score *χ*, defined as

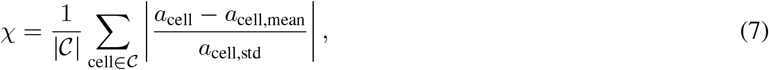

where 𝒞 is the set of neurons used, *a*_cell_ is the activity of a cell, and *a*_cell,mean_, *a*_cell,std_ are the mean and standard deviation of the activity of the same cell in the baseline simulation. Then, we selected ommatidia where *χ* is greater than a threshold *χ*_thr_ = 7 as the object. Once the object mask was detected, we calculated a descending signal using the method described in **“Visual object tracking task”** with the range of DN drives [DN_min_, DN_max_] set to [0.4, 1.2]. This descending signal was then passed to the hybrid walking controller to navigate either flat or “blocks” (height reduced to 0.2 mm) terrain.

In the schematic diagram **Figure 6e**, the placement of Tm and TmY neurons is based on [74]; the placement of T1 neurons is based on [75]; the placement of T2 and T3 neurons is based on [76]; the placement of T4 and T5 neurons is based on [77].

### Software

We used the following software in this study: Python 3.12, NumPy 1.26.4, SciPy 1.13.0, OpenCV-Python 4.9.0.80, Numba 0.59.1, Pandas 2.2.2 for general computing; Gymnasium 0.29.1, MuJoCo 3.1.4, dm control 1.0.18, PhiFlow 2.5.3 for physics simulation; PyTorch 2.2.2/2.3.0, PyTorch Lightning 2.2.2, PyTorch Geometric 2.5.0 for neural networks; Stable Baselines 3 2.3 for reinforcement learning; Nvidia graphics driver 550.54.15, Nvidia CUDA Toolkit 12.4 for GPU acceleration; FlyVision commit 056e4aa for connectome-constrained visual system simulation; SeqIKPy 1.0.0 for inverse kinematics; and Blender 2.81 for rigging the biomechanical model. Installation is managed automatically by package installers such as pip based on the `setup.py` file of our FlyGym package (see Code Availability statement).

## Data availability

Data are available via The Harvard Dataverse Repository at https://doi.org/10.7910/DVN/3MCEYR. This repository includes (i) the experimentally recorded walking kinematics, (ii) trained parameters of the path integration models, (iii) trained parameters of the head stabilization models, (iv) trained parameters of the visual processing and reinforcement learning models in the multi-modal navigation task, (v) training data for the visual processing model, and the graph representation of the ommatidia lattice used to perform graph convolution, (vi) the simulated complex plume dataset, and (vii) baseline neuron activities in the connectome-constrained visual system model.

## Code availability

The FlyGym package is available at https://github.com/NeLy-EPFL/flygym under the Apache-2.0 license. The documentation for FlyGym, along with detailed tutorials for some experiments in this paper, is available at https://neuromechfly.org.

The code used to generate some figures is not a part of the FlyGym package but is instead available at https://github.com/NeLy-EPFL/nmf2-paper under the same license.

A frozen snapshot of our code is available via Zenodo at https://zenodo.org/doi/10.5281/zenodo.12973000. However, FlyGym is under continued development and we recommend always using the latest version. Additionally, note that the results might not be bit-for-bit identical to the ones shown in this paper even with an exact copy of the code and its dependencies. This is due to differences in the computing hardware.

## Extended data figures

**Extended Data Figure 1:**
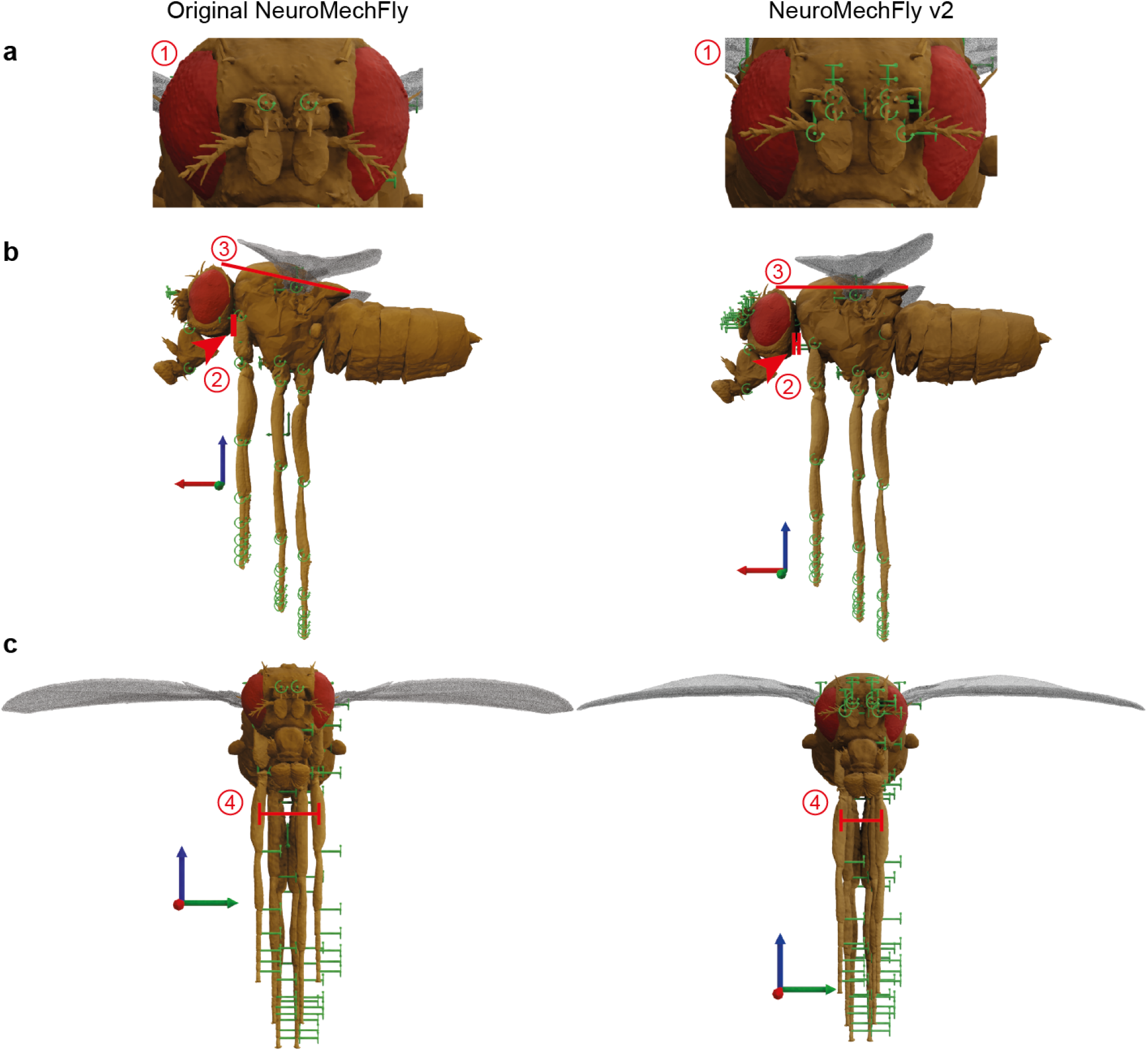
Improvements to the biomechanical model. A comparison of the original (left) and updated (right) NeuroMechFly biomechanical model from a **(a)** zoomed-in view of the head, highlighting antennal DoFs, **(b)** the side views, and **(c)** the front views. DoFs are indicated in green. The highlighted differences are: (1) additional DoFs in the antennae, (2) a gap for the neck between the head and the thorax, (3) angles of the thorax and the position of the head relative to it, and (4) the placements of the legs on the thorax.

**Extended Data Figure 2:**
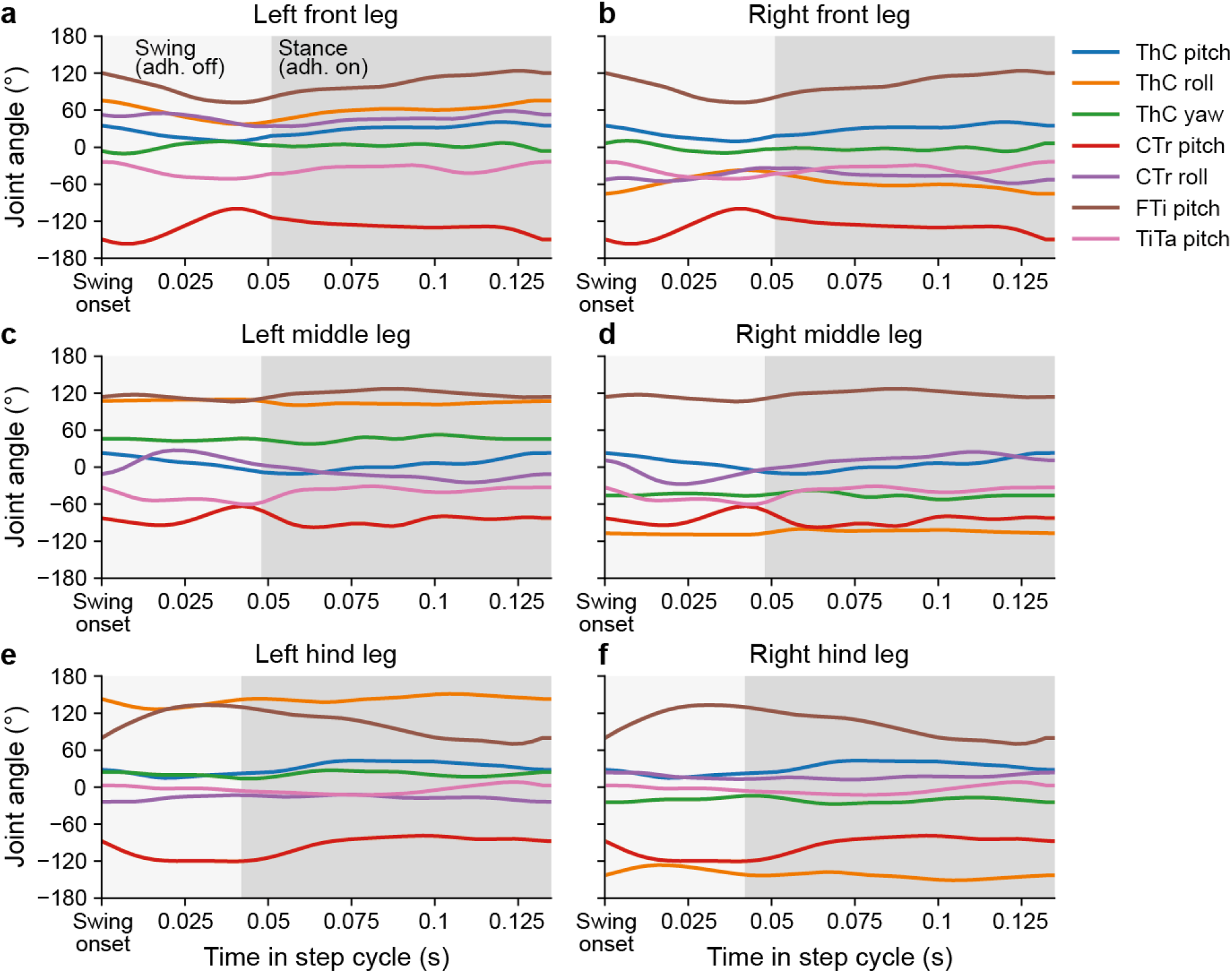
Preprogrammed stepping based on experimentally recorded data. Joint kinematics for each leg during preprogrammed stepping. Kinematic patterns derived from behavioral recordings. Time series for each joint are colorcoded. ThC: thorax-coxa joint; CTr: coxa-trochanter joint; FTi: femur-tibia joint; TiTa: tibia-tarsus joint. Note the left-right symmetry in roll and yaw DoFs. Periods when adhesion is turned off during swing to facilitate lifting each leg are indicated in light gray; periods when adhesion is on are indicated in dark gray.

**Extended Data Figure 3:**
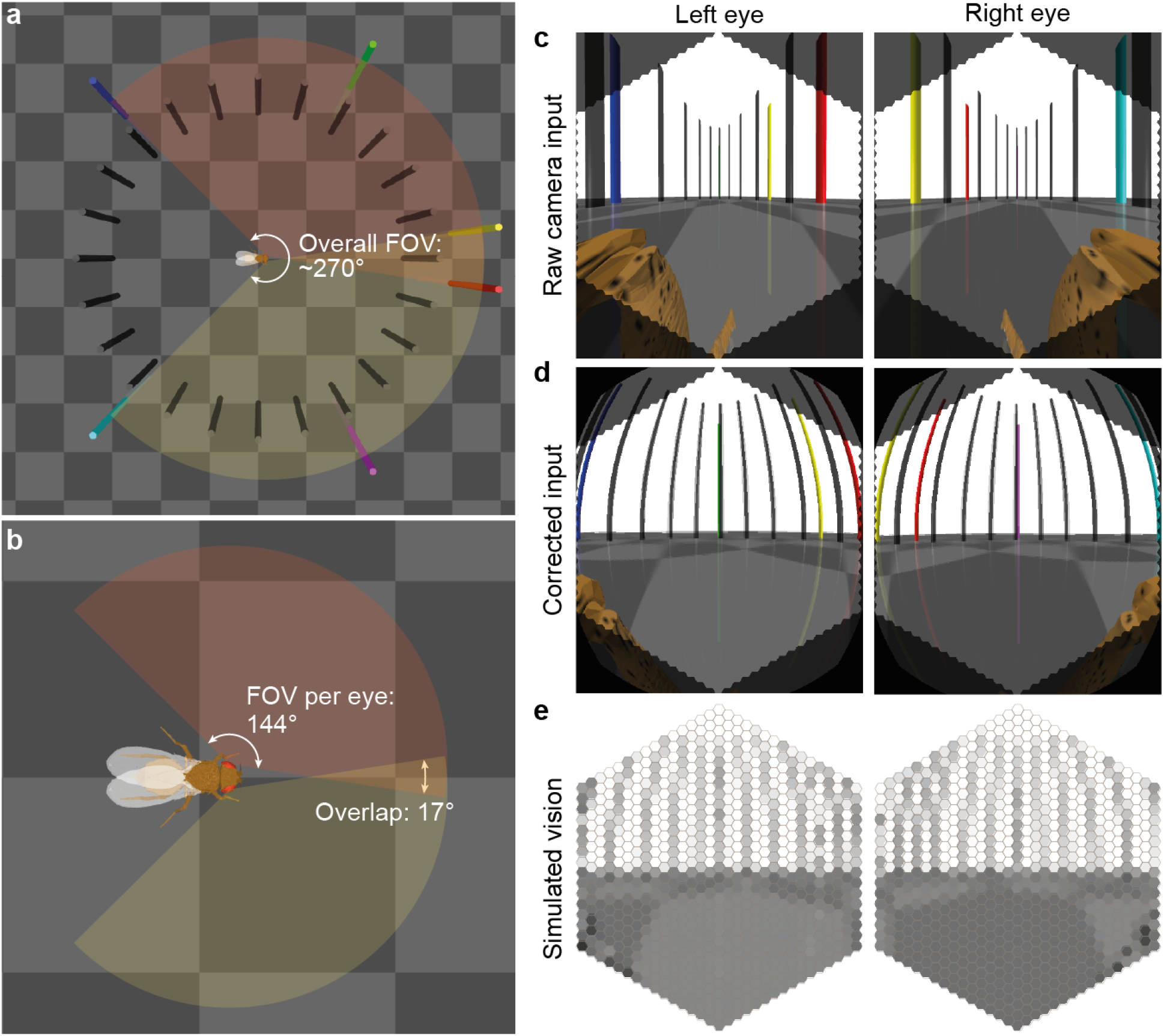
Calibration of vision. **(a)** The calibration environment has black pillars spaced regularly around the fly at 15 ° intervals. Additionally, red, green, and blue pillars are used to indicate the anterior, midline, and posterior field of view (FOV) limits of the left eye. Yellow, magenta, and cyan pillars indicate the FOV limits of the right eye. **(b)** Each eye has a FOV spanning ∼144 ° horizontally. The two eyes overlap by ∼17 °, resulting in an overall horizontal FOV of ∼270 °. **(c)** A raw camera view of what the fly sees in this environment before applying a fisheye effect. Note that by default, the rectilinear camera distorted areas closer to the edges of the FOV to keep the lines straight. **(d)** A fisheye effect is applied to simulate the roughly spherical arrangement of ommatidia in the fly eye. **(e)** Retinal inputs are simulated by binning the pixels according to the hexagonal grid of ommatidia and taking the average intensity within each ommatidium. Ommatidia are randomly sensitive to green (yellow-type) and blue (pale-type) channels in a 7:3 ratio.

**Extended Data Figure 4:**
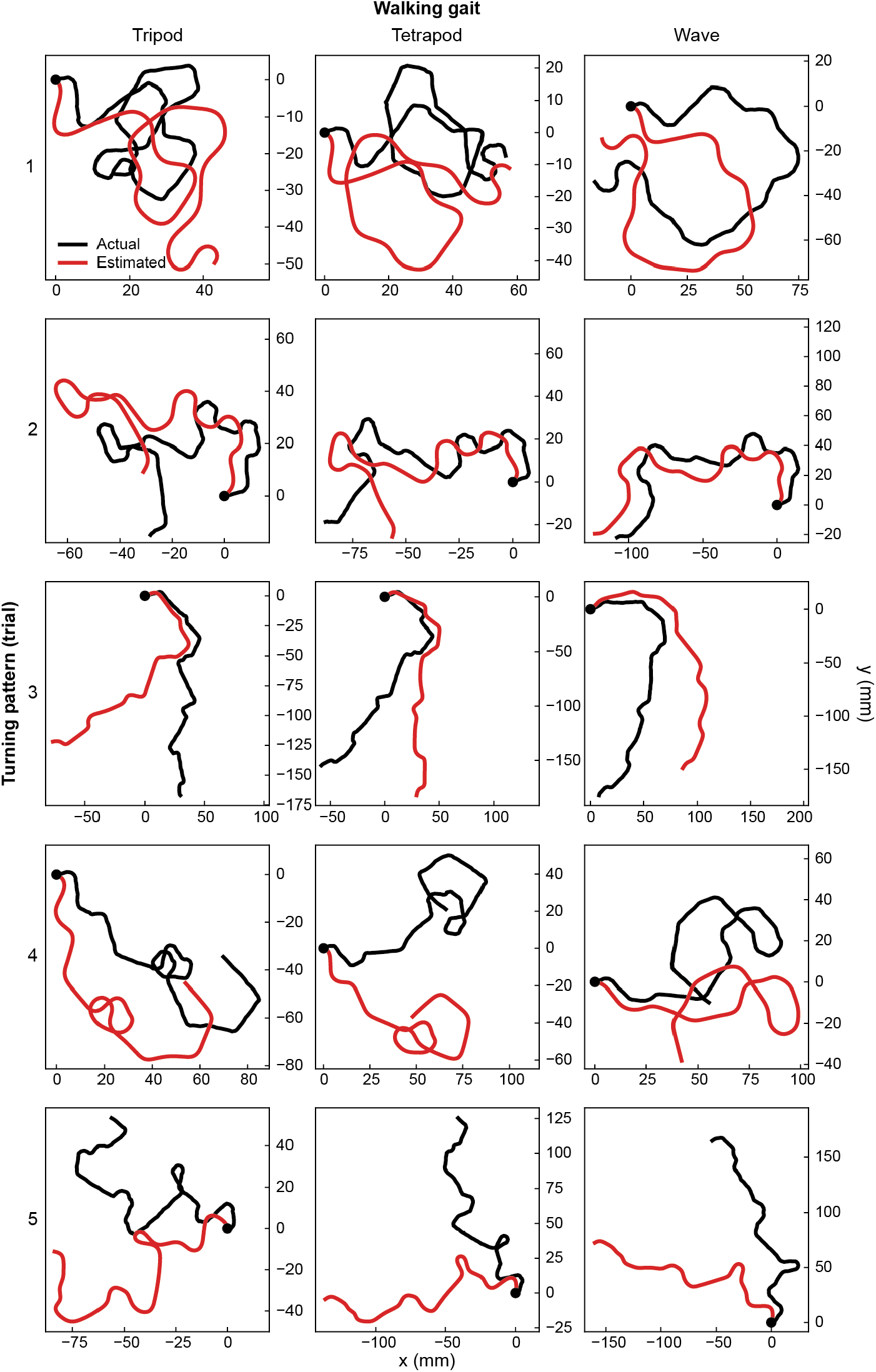
Trajectories during path integration based on ascending feedback. Actual (black) and ascending feedback-based (red) estimates of walking trajectories for five trials (rows) and three different insect locomotor gaits (columns). Indicated are starting positions of the paths (black circles). For each trial (row), the fly executes the same sequence of straight walking and turns but with different gaits.

**Extended Data Figure 5:**
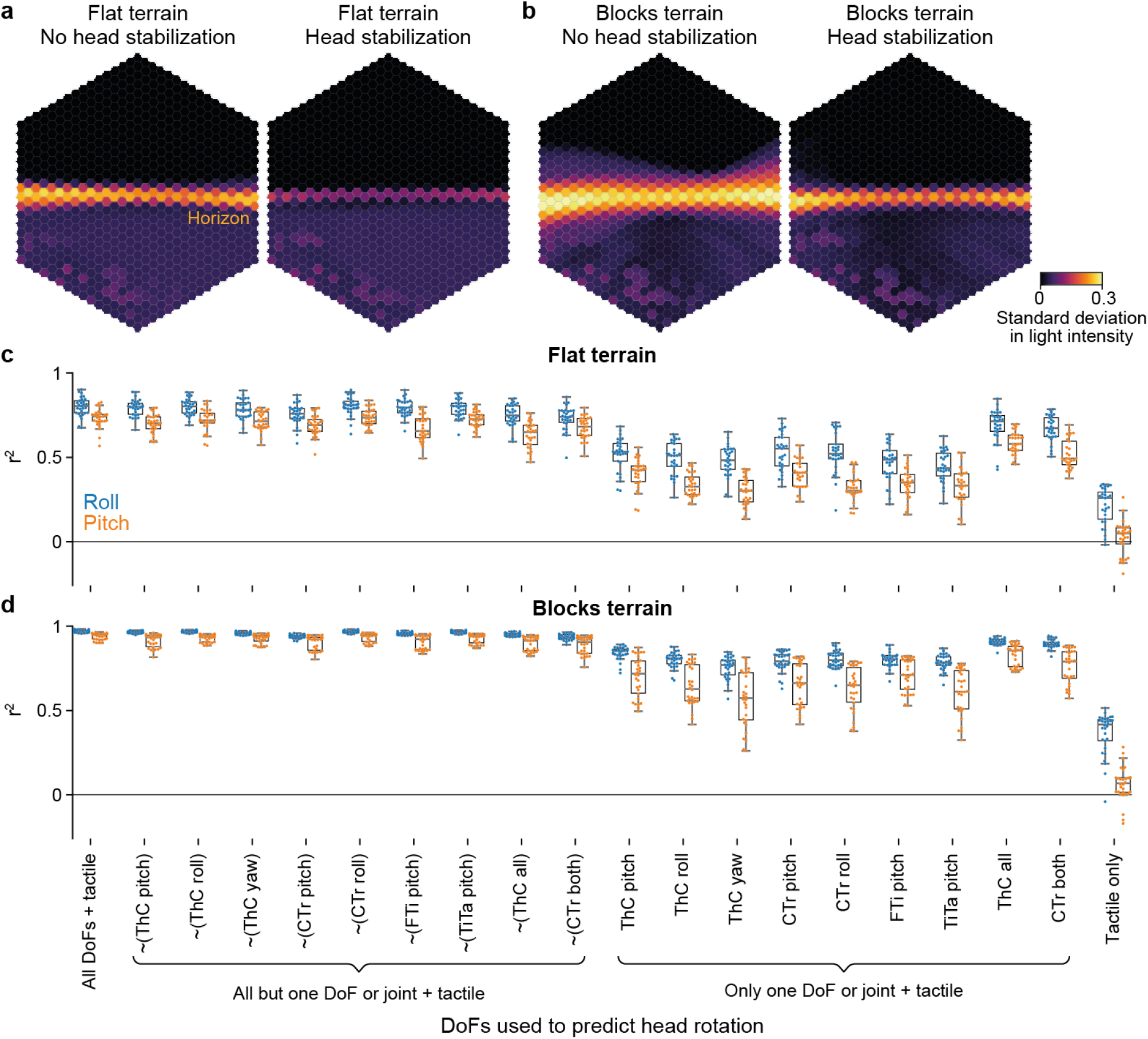
Efficacy of head stabilization as a function of terrain type and ascending signals. **(a–b)** The standard deviations of ommatidia readings from the left eye while walking over **(a)** flat or **(b)** blocks terrain without or with ascending feedback-based head stabilization. Note the high variability in light intensity near the horizon when the head is not stabilized; this is due to more pronounced self motion of the head. **(c–d)** Coefficient of determination (*r*^2^) between predicted and optimal head roll (blue) and pitch (brown) when performing head stabilization while walking over **(c)** flat or **(d)** blocks terrain and using ascending motor feedback from different sets of leg joint angles. “ ∼(removed DoF)” indicates all leg DoFs are used except the removed one (or ones at the same joint); “used DoF” indicates only the indicated DoF (or multiple ones at the same joint) are used. In each case, the same set of DoFs is used in all legs. Note that ground contact information is always used, hence the better-than-chance performance in the cases where no leg DoF is used. Overlaid are box plots indicating the median, upper and lower quartiles, and whiskers extending to the furthest points excluding outliers that are more than 1.5× the interquartile range (IQR) beyond the IQR. N=30 for each box; trials where physics simulation failed due to numerical instabilities are excluded.

**Extended Data Figure 6:**
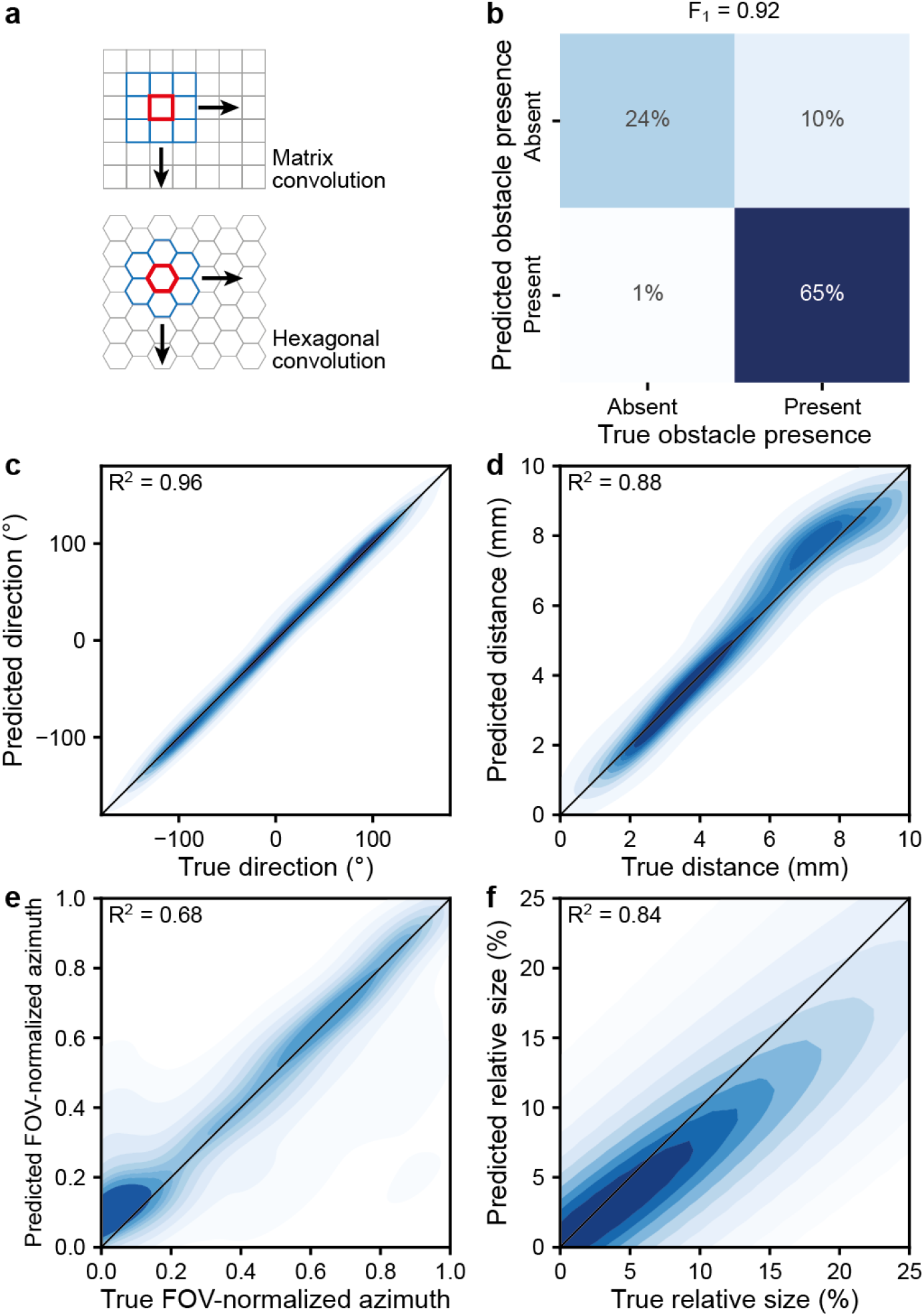
Vision model used in the multi-modal navigation task. **(a)** Illustration of the hexagonal convolution kernel compared to the standard matrix convolution kernel. **(b)** Accuracy of the model in predicting whether the obstacle is present in the fields of view of the fly’s eyes. The reported F1 score is the harmonic mean of the precision and recall. **(c–d)** Accuracy of the model in predicting the **(c)** direction and **(d)** distance of the obstacle from the fly. The angular *r*^2^ score is defined as the *r*^2^ score of sin(*ϑ*) concatenated with cos(*ϑ*) for all samples, where is the angle. **(e–f)** Accuracy of the model in predicting the **(e)** azimuth and **(f)** size of the obstacle in the retinal images. N=2,646.

**Extended Data Figure 7:**
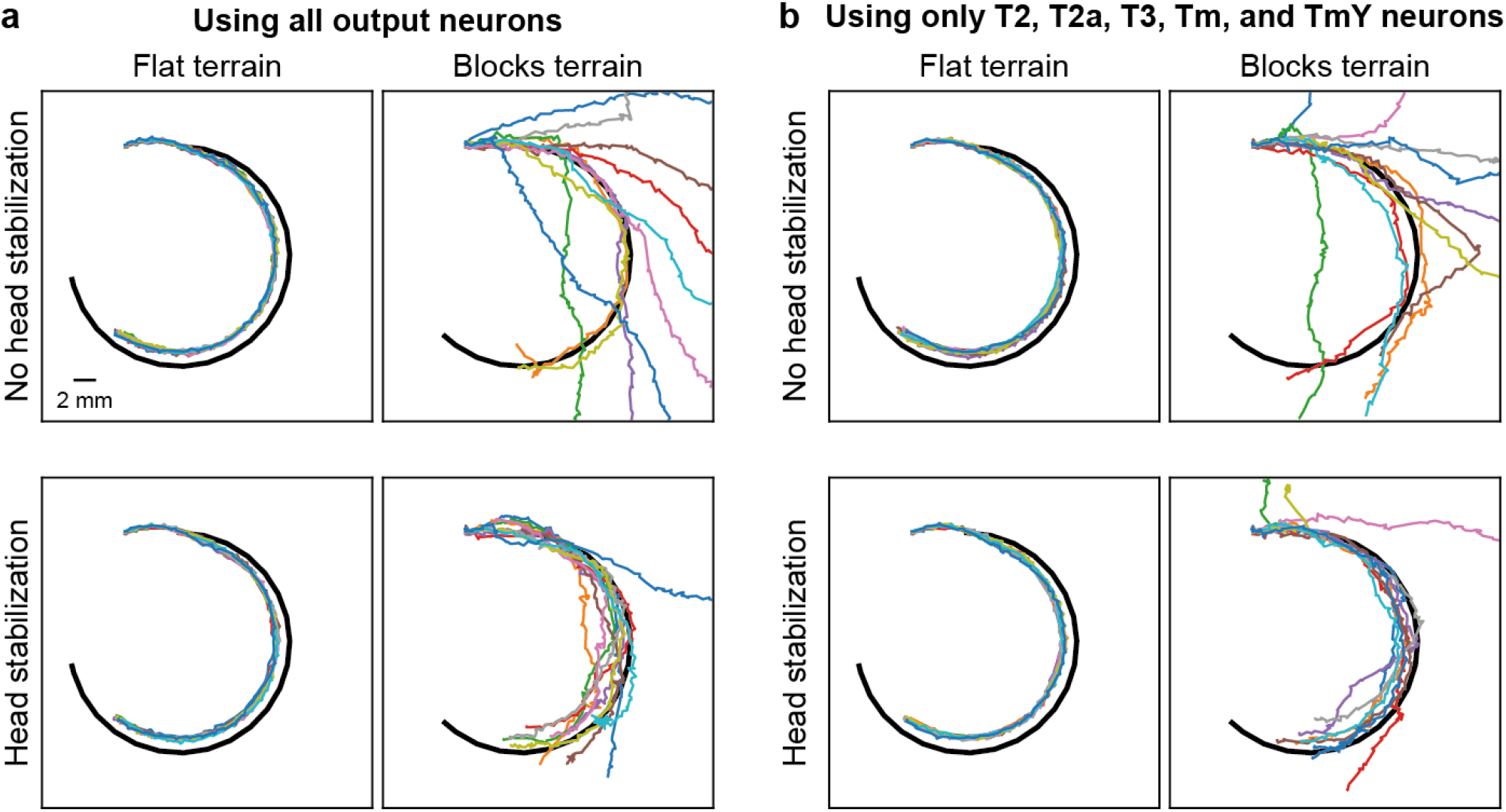
Performance of the connectome-constrained visual controller in a fly following task. Using **(a)** all T1–T5, Tm, and TmY neurons, or **(b)** T2, T2a, T3, Tm, and TmY neurons—upstream partners of LC9 and LC10 neurons [52]— to perform fly following either without or with head stabilization while walking over flat or blocks terrain. Shown are 11 trials (color-coded) per case.

## Videos

**Video 1: Obtaining 3D poses from an untethered walking fly to model more realistic stepping**. (Left) Comparison between template aligned (solid lines) and forward kinematics-reconstructed (dashed lines) 3D poses. Legs are colorcoded. (Right) Recording of an untethered fly walking straight through a corridor as seen from three viewpoints. Note that the center panel of the fly recording shows the ventral view; therefore, the legs closer to the top of the screen are on the left side of the fly. Video was recorded at 360 FPS, downsampled to 120 FPS, and displayed at 10% speed (i.e., 12 FPS). Overlaid are manually annotated thoracic, antennal, head, abdominal, and leg keypoints for 36 frames. Data used for pre-programmed steps are indicated (red lines).

URL: https://github.com/NeLy-EPFL/nmf2-videos/raw/main/Video1.mp4

**Video 2: Pre-programmed stepping pattern of each leg**. Individual legs are stepped in series according to their 3D pose estimation-derived joint kinematics. Simulation is played back at 0.05*×* real speed.

URL: https://github.com/NeLy-EPFL/nmf2-videos/raw/main/Video2.mp4

**Video 3: Ground reaction forces during locomotion with a CPG-based controller**. Here and in all subsequent videos, the tarsi are color-coded: natural leg color indicates that adhesion is off; dark blue indicates that adhesion is on and the tarsus is in contact with the ground; red indicates that adhesion is on but the tarsus is not in contact with the ground. In this particular video, because the terrain is flat, the third case rarely occurs. Simulation is played back at 0.05*×* real speed. URL: https://github.com/NeLy-EPFL/nmf2-videos/raw/main/Video3.mp4

**Video 4: Locomotion over sloped, vertical, and inverted terrain using leg adhesion**. Locomotion is driven by a CPG-based controller. Shown are simulations without (left) or with (right) leg adhesion. Indicated is the slope of the terrain (top). Simulation is played back at 0.1*×* real speed.

URL: https://github.com/NeLy-EPFL/nmf2-videos/raw/main/Video4.mp4

**Video 5: Control signals of the CPG-based controller**. Shown for all legs are the CPG phases (wrapped by 2*π*) and amplitudes from random initializations. As CPGs synchronize, they generate a tripod gait. Simulation is played back at 0.1*×* real speed.

URL: https://github.com/NeLy-EPFL/nmf2-videos/raw/main/Video5.mp4

**Video 6: Control signals of the rule-based controller**. Shown for all legs are the stepping scores and contributions of each of the three coordination rules. Indicated is the initiation of steps (triangles). Simulation is played back at 0.1 *×* real speed.

URL: https://github.com/NeLy-EPFL/nmf2-videos/raw/main/Video6.mp4

**Video 7: Control signals of the hybrid controller**. Shown for all legs are the CPG phases (wrapped by 2*π*) and amplitudes as well as the activation of the overstretch (solid) and stumbling (dashed) rules based on sensory feedback. The tibia is colored pink when the overstretch rule is active and light blue when the stumbling rule is active. Simulation is played back at 0.1*×* real speed.

URL: https://github.com/NeLy-EPFL/nmf2-videos/raw/main/Video7.mp4

**Video 8: Locomotion over multiple terrain types**. The fly walks over a flat surface (first column), a surface with gaps (second column), a surface with blocks (third column), and a mixed surface (fourth column). The fly is either controlled by a CPG-based controller (top), a rule-based controller (middle), or a hybrid controller which integrates both CPGs and sensory feedback rules (bottom). Shown are the results from 20 trials for each condition. For the hybrid controller, the tibia is colored pink when the overstretch rule is active and light blue when the stumbling rule is active. Simulation is played back at 0.1*×* real speed.

URL: https://github.com/NeLy-EPFL/nmf2-videos/raw/main/Video8.mp4

**Video 9: Visual object tracking task**. The fly uses vision to follow a black sphere that is moving away along an S-shaped trajectory. Shown are raw visual signals from the left and right eyes (bottom). A hybrid controller with leg adhesion is used for locomotion. Note that each eye’s field of view can observe front leg movements. Simulation is played back at 0.5 *×* real speed.

URL: https://github.com/NeLy-EPFL/nmf2-videos/raw/main/Video9.mp4

**Video 10: Olfactory chemotaxis task**. The fly seeks an attractive odor source (orange) while avoiding two aversive odor sources (blue). Colored bars (bottom) indicate the intensities of attractive (orange) and aversive (blue) odors sensed by antennae on each side of the head. A hybrid controller with leg adhesion is used for locomotion. Simulation is played back at 0.5*×* real speed.

URL: https://github.com/NeLy-EPFL/nmf2-videos/raw/main/Video10.mp4

**Video 11: Head stabilization using ascending motor feedback**. Shown are an overhead view of the fly, a zoom-in view of head movements, the raw left eye ommatidia signals, and time-series of neck actuation (roll and pitch) in the absence or presence of head stabilization. Neck actuation signals are either optimal (based on inverting thoracic pitch and roll, dashed lines) or predicted based on ascending motor feedback signals from the legs (solid lines). The first half of the video is during walking over flat terrain. The second half is during walking over blocks terrain. Simulation is played back at 0.2*×* real speed.

URL: https://github.com/NeLy-EPFL/nmf2-videos/raw/main/Video11.mp4

**Video 12: Neural controller for multimodal navigation trained through reinforcement learning**. The fly seeks an attractive odor source (orange) while using vision to avoid an obstacle (black pillar) over rugged mixed terrain. Shown are visual inputs to the left and right eyes (bottom-center). Orange bars (bottom-left and bottom-right) indicate the intensities of an attractive (orange) odor sensed by antennae on each side of the head. Locomotion is regulated using a hybrid controller with leg adhesion. Simulation is played back at 0.2*×* real speed. Nine trials beginning from different spawn locations are shown sequentially.

URL: https://github.com/NeLy-EPFL/nmf2-videos/raw/main/Video12.mp4

**Video 13: Navigating a complex plume using a bio-inspired odor-taxis algorithm**. The fly model uses a *Drosophila* plume navigation algorithm [51] to reach the odor source (left). Shown are the fly’s current state (e.g., pause, turn, walk forward) and algorithm parameter values (bottom-left). Red bar (bottom-left) indicates the intensity of odor detection. Also shown are the trajectory of the fly (red) and a zoomed-in birds-eye view of the fly (bottom right). Simulation is played back at 0.5*×* real speed.

URL: https://github.com/NeLy-EPFL/nmf2-videos/raw/main/Video13.mp4

**Video 14: Following another fly using a connectome-constrained vision network**. A “following” fly model controlled by a connectome-constrained visual system neural network [31], ascending motor feedback-based gaze stabilization, descending steering, and a hybrid locomotor controller with leg adhesion is tasked to follow a “leading” fly model across blocks terrain. Shown are an overhead view of both models’ trajectories (top row, middle column), raw ommatidia readings from the left and right retinas (top row, first and last columns), object detection scores obtained by processing visual neuron outputs (top row, second and fourth columns), and the spatial activities of *Drosophila* neurons processing visual signals from each eye (bottom). Neural polarization is color-coded from most hyperpolarized (blue) to most depolarized (red). Simulation is played back at 0.2*×* real speed.

URL: https://github.com/NeLy-EPFL/nmf2-videos/raw/main/Video14.mp4

## Supplementary Information

## 1 Supplementary Notes

### Note 1: Summary of supported features, demonstrated use cases, and future directions

This note summarizes features generally supported by the NeuroMechFly framework, modeling choices specifically demonstrated in this paper, and directions for future work.

#### Visual system

- Currently supported:

- Raw retina readings processed by any user-defined visual system model

- Shown in examples:

- Object size and location on the retina
- Artificial neural network (convolution on a hexagonal lattice)
- Connectome- and physiology-constrained visual system model [1]

- Future potential:

- Algorithmic visual system model [2]
- Robotics/AI: Vision models at variable levels of abstraction

#### Olfactory system

- Currently supported:

- Intensity readout in an n-dimensional space according to a user-defined diffusion relationship
- User can decide where sensors are placed on the antennae and/or maxillary palps
- Signals can be processed with any user-defined olfactory system model

- Shown in examples:

- Readout of 2D (attractive and aversive) odor intensities by antennae and maxillary palps
- Simple model computing the difference in odor intensities between bilateral olfactory organs
- Model of complex odor plumes
- Algorithmic odor plume processing model [3]

- Future potential:

- Simulation of fluid interaction with the animal’s body and its movements [4, 5].
- Alternative algorithmic odor plume processing models [6, 7, 8]
- Connectome- and physiology-constrained olfactory system models [9]
- Robotics/AI: Odor search models at variable levels of abstraction

#### Action selection model

- Currently supported:

- Arbitrary user-defined process with access to visual/olfactory signals, ascending motor signals, and internal states

- Shown in examples:

- Steering based on visual or olfactory object detection
- Behavior state transitions governed by Poisson processes
- Black-box multi-layer perceptron (MLP) trained by reinforcement learning

- Future potential:

- Algorithmic action selection models
- Connectome- and physiology-constrained action selection models [10, 11]
- Robotics/AI: Action selection models at variable levels of abstraction

#### Descending controller

- Currently supported:

- n-dimensional descending space defined by the user

- Shown in examples:

- 2D descending drive modulating Central Pattern Generators (CPGs) on each side of the body

- Future potential:

- High-dimensional descending representation allowing command-like and population-based coding [12]

#### VNC (motor control) model

- Currently supported:

- Arbitrary user-defined VNC model/process with access to mechanosensory signals, internal states, and descending signals

- Shown in examples:

- CPG-based controller [13]
- Sensory feedback rule-based controller [14]
- Hybrid controller integrating CPGs and sensory feedback rules

- Future potential:

- Additional algorithmic motor system models
- Connectome- and physiology-constrained motor system models [15, 16]
- Robotics/AI: Hexapod motor system models at variable levels of abstraction [17]

#### Ascending feedback

- Currently supported:

- n-dimensional ascending space defined by the user

- Shown in examples:

- Leg stride lengths for path integration
- Leg joint angles and ground contacts for head stabilization

- Future potential:

- High-dimensional ascending representations encoding behavioral and internal states at variable levels of abstraction [18]
- Robotics/AI: Ascending motor signals at variable levels of abstraction [19]

#### Behavioral kinematics

- Currently supported:

- User-defined limb kinematics, either estimated from recordings of real animals or generated by models
- User-defined sets of actuated versus passive degrees of freedom (DoFs)
- User-defined rules to turn adhesion on/off

- Shown in examples:

- Each walking step: 42 actuated leg degrees of freedom (DoFs, 7 per leg) playing out pre-programmed stepping sequences
- Stepping coordination: see walking controller model
- Adhesion on/off periods defined as a part of pre-programmed stepping sequences

- Future potential:

- Inclusion of motor neurons and muscle models [20, 21]
- Real-time replay of animal behavior [22], which enables closed-loop experimentation [23]
- Robotics/AI: Body movement patterns learned by imitating animals [24]

#### Environment

- Currently supported:

- User-defined arena with static (e.g., rugged terrain, obstacles, odor sources) or dynamic (e.g., moving objects, moving odor sources, plumes) features

- Shown in examples:

- Low-level features: Rugged terrains for benchmarking locomotor controllers
- Higher-order features: Odors, obstacles, and a moving sphere for sensory-guided locomotion, additional fly models for social behavior [25, 26]

- Future potential:

- Aerodynamics for flight and odor plume propagation
- Compliant objects
- Complex, visually realistic terrain

#### Parametrization/training algorithm

- Currently supported:

- Manual parameter tuning or arbitrary user-defined optimization algorithm (e.g., genetic algorithms, reinforcement learning algorithms)

- Shown in examples:

- Manual parameter tuning
- Gradient-based reinforcement learning algorithm (SAC [27])

- Future potential:

- Larger-scale simulation, optimization, and neural architecture search

#### Physics simulator

- Currently supported:

- PyBullet (original version [28])
- MuJoCo [29]

- Shown in examples:

- MuJoCo [29]

- Future potential:

- GPU-accelerated and/or gradient-aware simulators for large-scale model training [30, 31, 32, 33]

### Note 2: FlyGym control loop

#### Algorithm 1

The main control loop in FlyGym

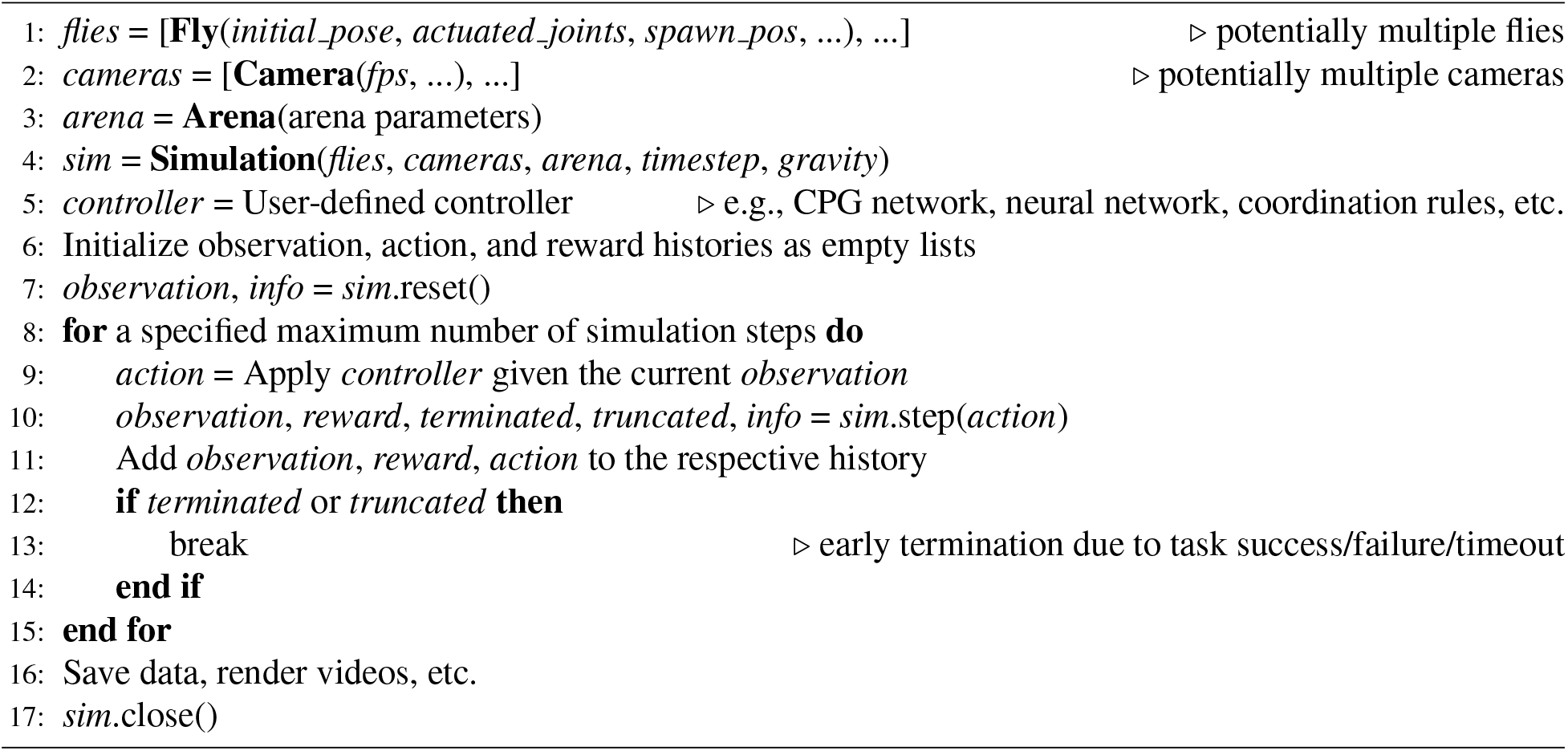

Algorithm 1 is a summary of the main control loop in our FlyGym library in pseudocode. The following is a breakdown of the steps:

- *Lines 1–4:* First, we define the simulated fly (or potentially a group of flies) by specifying the initial pose, actuated DoFs, spawn position, etc. Then, we define a list of cameras with parameters specifying how the simulation should be rendered. Next, we define the arena with parameters such as arena size. With these, we can define the simulation object itself, specifying parameters such as the time step size.
- *Line 5:* The implementation of the controller is determined by the user. Typically, it has a prediction function that determines the action to take given the current observation. An internal state can be maintained within the controller. The controller can be a hard-coded model or an artificial neural network (e.g., a network trained through reinforcement learning). The definition of the controller (or learning algorithm) is isolated from the control loop.
- *Lines 8–15:* This is the main Markov Decision Process (MDP) loop. Using the user-defined controller, we supply the appropriate action to the simulation and step it forward by one simulation time step. The simulation returns a new observation, optionally a reward (depending on the user’s definition), a “terminated” flag indicating whether the simulation has ended due to a factor within the MDP framing (e.g., task success or failure), a “truncated” flag indicating whether the simulation has ended due to a factor outside of the MDP framing (e.g., simulation time reached, physics error), and a dictionary containing any additional information.
- *Lines 16–17:* The user can then save the necessary data and end the simulation.

### Note 3: Action and observation spaces of the default Markov Decision Process

#### Action space

- *Joint actuation signals:* The target angle, angular velocity, or torque (as specified by the user) of the n actuated joint degrees of freedom (by default all 42 leg degrees of freedom (DoFs)), ℝ^*n*^.
- *Adhesion on/off signals (if adhesion is enabled):* Whether adhesion is turned on or off at the tip of each leg, *{*0, 1*}*^6^.

#### Observation space (for each fly)

- *Joint states:* The angle, velocity, and force at each of the n actuated joint DoFs (by default all 42 leg DoFs), ℝ^3*×n*^.
- *Fly state:* The linear (x-y-z) position, linear velocity, angular (pitch-roll-yaw) position, and angular velocity of the base of the fly (thorax) in the arena in the global reference frame, ℝ^4*×*3^.
- *Fly orientation:* Unit vector aligned with the anterior-posterior axis of the fly in the global reference frame, ℝ^3^.
- *Ground reaction forces*: The ground reaction forces in all three dimensions when experienced by the m body segments (by default the tibia and all 5 tarsal segments of all legs) in the global reference frame, ℝ^*×*3^.
- *Visual inputs (if vision is enabled):* Light intensity readings from all 721 ommatidia on each eye, ℝ^2*×*721*×*2^. Note that 70% of the ommatidia are yellow-type and 30% are pale-type. The last dimension indicates the color channel (for each ommatidium, only one of the two values is non-zero).
- *Odor inputs (if olfaction is enabled):* Odor intensities sensed by the antennae and maxillary palps on two sides of the fly in a k-dimensional odor space, ℝ^*k×*4^.

### Note 4: Network of Central Pattern Generators

The dynamics of the Central Pattern Generator (CPG) network are governed by the following differential equations:

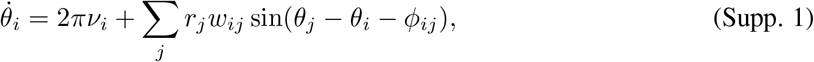

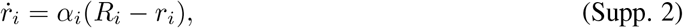

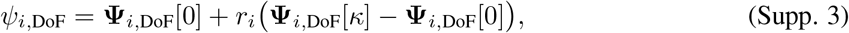

where *θ*_*i*_ and *r*_*i*_ are the state variables describing the phase and amplitude of the oscillator for leg *i, ν*_*i*_ and *R*_*i*_ are the intrinsic frequency and amplitude of leg *i, α*_*i*_ is a constant determining the rate of convergence to the intrinsic amplitude, *w*_*ij*_ and *ϕ*_*ij*_ are the coupling weight and phase bias between legs *i* and *j, ψ*_*i*,DoF_ is the angle of the degree of freedom DoF on leg *i*, **Ψ**_*i*,DoF_[*t*] gives the angle of DoF at time *t* of the preprogrammed step cycle, and **Ψ**_*i*,DoF_0] is the neutral position (start-of-swing pose). The index in the time series, *κ*, is then given by

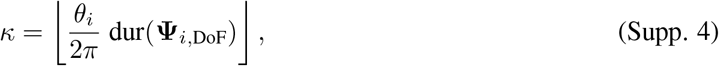

where dur(**Ψ**_*i*,DoF_) is the duration of the preprogrammed step.

### Note 5: Calibration of visual input

To calibrate vision, we surrounded the model with color-coded pillars in the arena **(Extended Data Figure 3)**. This allowed us to identify the limits and mid-lines of the field of view (FOV) of each eye (i.e., camera) and configure the camera’s orientation and FOV correctly. MuJoCo’s built-in camera renders a rectilinear image with a distortion: the periphery of the image is enlarged to keep lines straight. This distortion is especially severe at large FOV, such as those used to simulate fly vision. To remove this distortion, we applied a fisheye effect on the raw camera readings **(Extended Data Figure 3c–d)** to represent the same angular span approximately equally in the rendered image. More precisely, the pixel on row r, column c in the transformed image is assigned the value of the pixel on row *r*^⋆^, column *c*^⋆^ of the original image, where *r*^⋆^, *c*^⋆^ are determined by

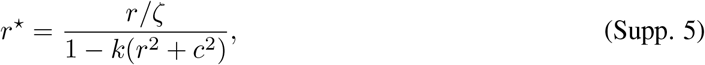

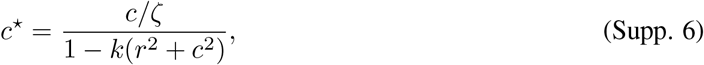

where *r, r*^⋆^, *c, c*^⋆^ are normalized to the range [− 1, 1] (i.e., the pixel at the center of the image has the coordinates 0, 0]); k = 3.8 describes the extent of the fisheye distortion, and *ζ* = 2.72 describes the zoom level of the resulting image. Where *r*^⋆^, *c*^⋆^ are outside the range [ −1, 1], an empty value is assigned. Finally, we binned the pixels by the ommatidia they belong to and took the mean intensity per ommatidium **(Extended Data 3e)**.

This process is accelerated with just-in-time compilation using Numba [34].

### Note 6: Path integration algorithm

We defined the ascending signals to be used for path integration such that they are limited to biologically plausible proprioceptive and tactile information. For each leg *L*, we calculated the positions of the claw relative to the body over time, **x**_*L*_(*t*). We also obtained a binary time series *p*_*L*_(*t*) indicating whether or not the leg is executing a power stroke (i.e., in stance). In our simulation, *p*_*L*_(*t*) = 1 if and only if the ground contact force of leg L is greater than a threshold *F*_thr_ in magnitude. We also tracked the heading of the fly over time, *h*(*t*). For convenience, we alternatively refer to the heading unit vector 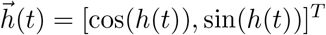 . This allowed us to calculate the projection of **x**_*L*_(*t*) in the direction of the fly’s current heading, 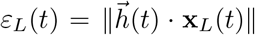 . Then, we integrate this proprioceptive signal, masked by *p*_*L*_(*t*), to obtain the cumulative stride length *d*_*L*_ for each leg. Intuitively, *− d*_*L*_ quantifies the accumulation of the body’s displacement with regard to the claw of leg L during stance in the direction of the fly’s orientation:

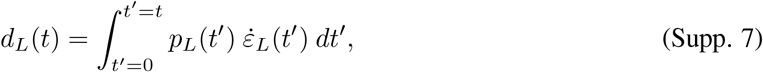

or, in discrete-time simulation where time steps are indexed by *i*,

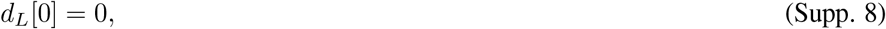

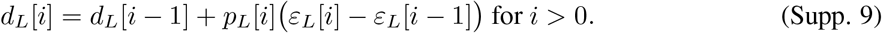

Next, we defined the left-right asymmetry (*δ*) and sum *(σ*) in this cumulative stride length for each pair of legs (using the continuous-time notation for simplicity):

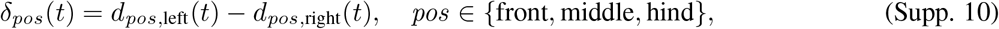

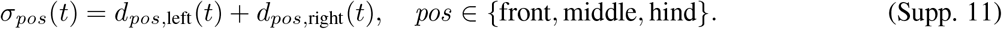

Intuitively, the change in *δ*_*pos*_ and *σ*_*pos*_ over time allows one to estimate the change in the fly’s heading and forward displacement, respectively.

Having defined the ascending mechanosensory signals, we then built regression models to estimate the change in body state. These models use the change in the cumulative ascending signals (defined above) over some time window *τ* to estimate the change in heading and cumulative forward displacement over the same time window. Note that for this purpose, different combinations of legs might be used for path integration (e.g., front legs only, front legs and middle legs, all legs). We defined our models as

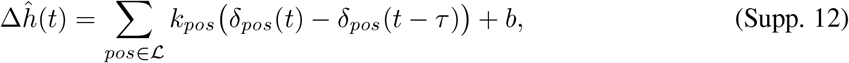

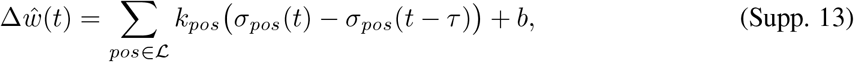

where ∆*ĥ*(*t*) and ∆*ŵ*(*t*) are the model-estimated change in heading and forward displacement over the past *τ* seconds; ℒ *∈* PowerSet( {front, middle, hind}) is the set of legs used for path integration; *k*_*pos*_, *b* are fitted parameters.

Then, we collected the real change in heading ∆*h*(*t*):

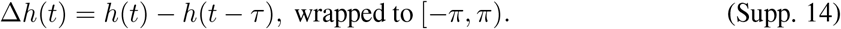

To obtain the real change in forward displacement ∆*w*(*t*), we need to first calculate the cumulative forward displacement *w*(*t*):

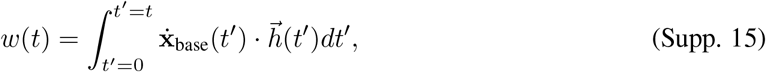

where **x**_base_ is the base position of the fly. In discrete time, this is

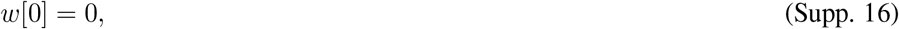

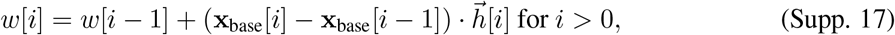

and, again using the continuous-time notation for simplicity,

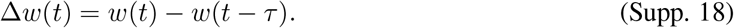

We then fit a set of models (for both heading and displacement), by least squares, for each of the first 10 trials of each gait type.

Once ∆*ĥ*(*t*) and ∆*ŵ*(*t*) are obtained through the model, we can perform path integration by essentially reversing the process described above. We obtain the model-estimated heading *ĥ*(*t*) and fly position 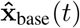 as follows:

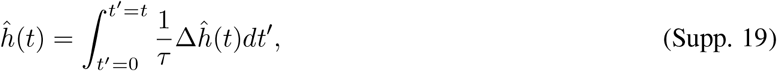

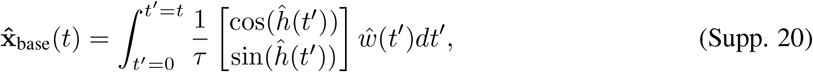

or, in discrete time,

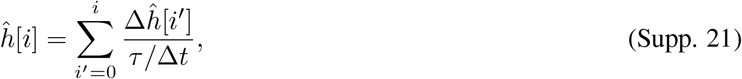

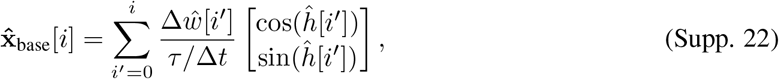

where ∆*t* is the simulation time step. Note that *ĥ* and 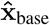 are defined only after *τ* seconds of simulation because proprioceptive information from *τ* seconds before the current time is needed to estimate the ∆*ĥ* and ∆*ŵ*. For simplicity, we start path integration only after *τ* seconds.

We chose *F*_thr,front_ = 0.5 mN, *F*_thr,middle_ = 1 mN, *F*_thr,hind_ = 3 mN, *τ* = 0.64 s through a sensitivity analysis and used all legs to reconstruct the path. We used an ensemble of the 10 trained models (by averaging parameters, since the model is linear) for prediction over the last 5 trials. The final path integration results reported in **Figure 4b–c** and **Extended Data Figure 4** are performed on the last 5 trials that were never used during training or parameter selection.

### Note 7: Head stabilization model

To establish the mapping from proprioceptive information to the appropriate neck movements, our strategy is to first simulate walking without control of neck joints, calculate the desired neck actuation signals based on the resulting head angle changes, and finally train a model to predict these neck movements using proprioceptive signals. A caveat is that by this approach, we consider the change in body inertia due to head movement, and therefore the impact on body physics negligible.

We next simulated walking over flat and “blocks” terrains (height of blocks reduced to 0.2 mm) using tripod, tetrapod, or wave gaits. We simulated both straight walking and turning using the hybrid turning controller. To sample different turns, we defined a scalar *b* ∈ [− 1, 1] which determines the direction and sharpness of the turn, with *b* = − 1, *b* = 0, and *b* = 1 indicating the sharpest turn to the left, straight walking, and the sharpest turn to the right, respectively. The descending drives for the inner and outer sides of the turn are defined as

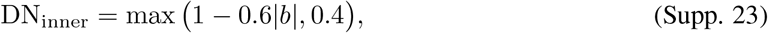

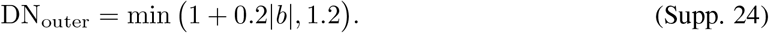

We then sampled different values of *b* to constitute the training-and-validation set (*b* ∈ {−1, −0.9, −0.8, …, 1}) and the out-of-sample testing set (*b* ∈ {−0.95, −0.85, −0.75, …, 0.95}). For each combination of terrain type, gait, and descending drive, we simulated 0.5 s of walking. The x-and y-coordinates of the fly’s spawn position are randomly sampled from a uniform distribution ranging from 0 to 1.3 mm, where 1.3 mm is the feature size of the repeating motif of the “blocks” terrain. We manually excluded simulations over “blocks” terrain in which the fly flipped.

Then, we extracted the input (mechanosensory information) and target output (ideal neck actuation) of the head stabilization model from the simulations. The input signals consist of the leg DoF angles and the leg contact forces. The angles of each leg DoF are standardized (i.e., subtracted by the mean and divided by the standard deviation) based on the angles distribution of the corresponding DoF during forward walking (i.e., *b* = 0) over flat terrain. Ground contact for each leg is calculated as the sum of the magnitudes of forces experienced by all tarsal segments, binarized using a threshold *F*_thr_. Consistent with the path integration task, we used *F*_thr,front_ = 0.5 mN, *F*_thr,middle_ = 1 mN, *F*_thr,hind_ = 3 mN for the three leg pairs. The target output signals are the optimal roll and pitch angles of the neck joint. To calculate these angles, we inverted the quaternion describing the 3D rotation of the thorax, converted the inverted rotation into Euler angles, and took the roll and pitch components. As we are not correcting for the yaw, this operation is independent of the orientation of the fly on the horizontal plane. Therefore, the model has a 48-dimensional input (7 DoF angles per leg *×* 6 legs + 6 contact variables) and a two-dimensional output (target neck roll and pitch).

With the model inputs and outputs extracted, we next trained a simple multi-layer perceptron (MLP) with two hidden layers, each with 32 units, with ReLU activation. Snapshots from the simulation are treated as independent samples. We trained the model by minimizing the mean squared error (MSE) using the Adam optimizer [35] at a learning rate *η* = 0.001 in batches of 256 samples. The training-and-validation set is further split into a training set and a validation set at a ratio of 8:2, each sampling from all available values of *b*. Using PyTorch [36] and Torch Lightning [37], we trained the model for 30 epochs on the training set and selected the best parameters based on the performance on the validation set. A single model is trained to handle all three gait types over both flat and “blocks” terrains. The *r*^2^ scores of the model in each simulation from the testing set are reported for statistics. Finally, we retrained the model using different combinations of joint DoFs in the inputs. The excluded inputs are set to 0 instead of removed so that the model architecture remains fixed for all variants.

To deploy the model in closed loop, we calculated the model-predicted neck angles and applied them using a proportional derivative (PD) controller on the joint angle with a gain of 500. Note that due to the naming convention chosen in the original NeuroMechFly model [28], the roll DoF of the neck is actually named `joint Head yaw` in the biomechanical model definition. For consistency, we decided to highlight this issue in the documentation of the FlyGym library but not change the DoF name.

### Note 8: Visual feature extraction model

We trained a visual feature extraction model that performs convolution on the retinal image. Because the retinal image is on a hexagonal lattice, we built a graph representation of the lattice and performed graph convolution [38] on it **(Extended Data Figure 6a)** using PyTorch [36] and PyTorch Geometric [39]. The model comprises the following layers. Note that in a later experiment, we used a more realistic, connectome-constrained visual system model.

1. Graph convolution layer (GCNConv): 2 in channels, 4 out channels
2. Hyperbolic tangent activation function (Tanh)
3. Graph convolution layer (GCNConv): 4 in channels, 2 out channels
4. Hyperbolic tangent activation function (Tanh)
5. Concatenation of left/right outputs
6. Linear layer: 2 *×* 2 *×*721 = 2, 884 inputs, 16 outputs
7. Hyperbolic tangent activation function (Tanh)
8. Linear layer: 16 inputs, 16 outputs
9. Hyperbolic tangent activation function (Tanh)
10. Linear layer: 16 inputs, 7 outputs

Given the ommatidia intensity readings as inputs, this model predicts (i) the direction of the object relative to the fly, (ii) the distance of the object from the fly, (iii) whether the object is within the fly’s eyes FOVs, (iv) the azimuth of the object seen from each eye, and (v) the percentage size of the object on the retina. We gathered 10,000 training samples by randomly spawning the fly and the obstacle (within 10 mm of the fly) and injecting Gaussian noise (standard deviation = 0.05 *×* the intensity range) into the visual inputs. The labels are calculated similarly to the approach used for simple visual tracking **(Figure 3a–c)** at an intensity threshold of 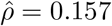. We then trained the model in a supervised manner using the Adam optimizer [35], at a learning rate of *η* = 0.001, with the loss *L* defined as

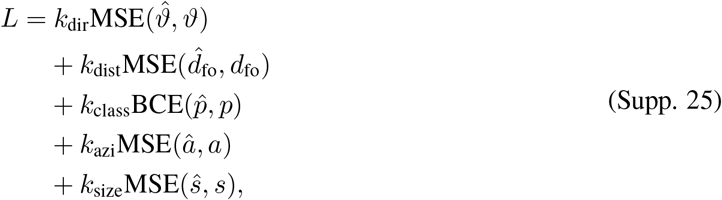

where *k*_dir_, *k*_dist_, *k*_class_, *k*_azi_, *k*_size_ are the coefficients for the obstacle direction, obstacle distance, obstacle presence classification, obstacle azimuth, and obstacle size terms; MSE is the Mean Squared Error, BCE is the Binary Cross Entropy; *ϑ* is the direction of the obstacle from the fly; *d*_fo_ is the distance of the obstacle from the fly; *p* is the predicted probability on whether the obstacle is within the fields of view of the fly’s eyes; *a* is the azimuth of the obstacle on the retina from each eye; *s* is the relative size of the obstacle on the retina from each eye; 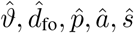 are the model’s predictions of *ϑ, d*_*fo*_, *p, a, s*. We empirically chose the following loss weights: *k*_dir_ = 0.5, *k*_dist_ = 2, *k*_class_ = 0.1, *k*_azi_ = 4, *k*_size_ = 2. We obtained satisfactory model performance after 150 epochs of training (**Extended Data Figure 6**).

### Note 9: Reinforcement learning task

The multi-modal navigation controller receives as inputs (i) the visual features preprocessed by the convolutional neural network described in **Supplementary Note 8**, and (ii) preprocessed olfactory information. To derive the preprocessed olfactory features, we calculated the odor intensity sensed on each side of the head by averaging the intensities sensed by the ipsilateral antenna and maxillary palp at a relative weight of 9:1. We then transformed the odor intensity with the square root operator to give it a more linear response as a function of distance. We concatenated the odor intensities on each side of the head with the extracted visual features and the turning bias *b* from the previous time step (described below). This feature vector was given to the controller as input.

The controller is a multi-layer perceptron (MLP) with 2 hidden layers, each with 32 units. It predicts a scalar turning bias *b*, which is converted to a descending representation according to **Equation 5** in the Methods. The turning bias is updated every 0.05 s of simulated time. We train this MLP using reinforcement learning: If the fly has reached within 3 mm of the target, a reward of *R*_success_ = 10 is given and the simulation is terminated. If the fly has flipped or touched the obstacle, a reward of *R*_fail_ = *−*5 is given and the simulation is terminated. If none of these special conditions are met, we calculate a score S_obstacle_ describing whether the fly is facing the obstacle: *S*_obstacle_ = 1 when *ϑ* = 0; *S*_obstacle_ = 0 when *ϑ* > *m* · arctan(*r*/*d*_fo_) where *r* is the radius of the obstacle; *d*_fo_ is the fly-to-obstacle distance. In other words, this score is 1 when the fly is perfectly facing the obstacle and 0 when the fly is facing away from the obstacle by a margin of *m* mm. When the fly’s heading is within this range, *S*_obstacle_ is linearly interpolated. Similarly, we calculated a second score *S*_target_ that is 1 when the fly is facing any point within *ϵ* mm of the target; by contrast, *S*_target_ = 0 when the shortest distance from the fly’s heading direction to the target is more than *qϵ. S*_target_ is also linearly interpolated when the shortest distance from the target to the fly’s heading vector is between *ϵ* and q*ϵ*. We empirically chose the following parameters: *m* = 4 mm, *ϵ* = 2 mm, *q* = 3. Then, we calculated the reward *R* defined as

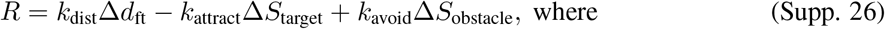

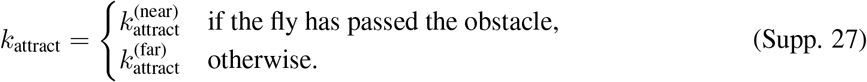

*k*_dist_, *k*_attract_, and *k*_avoid_ are the weights of the proximity, target facing, and obstacle-avoiding terms of the reward. *k*_attract_ has two possible values depending on the stage of the task. *d*_ft_ is the distance from the fly to the target. ∆*d*_ft_, ∆*S*_target_, and ∆*S*_obstacle_ are the current values of *d*_ft_, *S*_target_, and *S*_obstacle_ subtracted from their values in the previous decision-making time step. We chose the following weights for the reward function: *k*_prox_ = 1, *k*_avoid_ = 7, 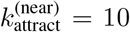, and 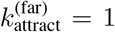. We trained the model using the Soft Actor-Critic (SAC) algorithm [40] implemented in the Stable Baselines 3 library [41], which uses gradient back-propagation to update the weights and biases of the MLP. We used the Adam optimizer [35] at a learning rate of *η* = 0.01. We trained the model for 500,000 decision steps (approximately 6.9 simulated hours). We consider this training time reasonable as the network is randomly initialized and the simulated fly has to “evolve” the entire circuit for decision-making.

### Note 10: Complex plume simulation and tracking

We modeled the plume as a distinct species embedded in an incompressible fluid. This is analogous to, for example, food odor being embedded in the air. As such, we simulated the plume by solving the Navier–Stokes equations in 2D:

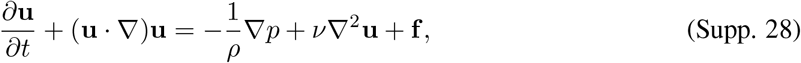

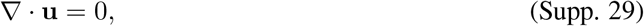

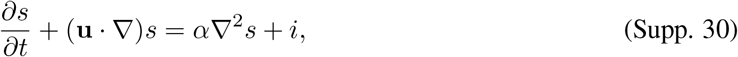

where **u** is the 2D velocity vector, *ρ* = 1 is density, *p* is the pressure, *ν* is the viscosity of the fluid, **f** is the external force vector, *s* is plume concentration, *t* is time, *α* is the diffusivity of the plume, and *i* is the inflow of the plume. In our simulation, *i* is 0.2 at an odor source with a radius of 2 mm centered at *x* = 10 mm, *y* = 80 mm and 0 everywhere else. Our simulation is initialized with a constant velocity field **u** = [1, 0]^*T*^, producing a left-bound “wind.” **f** is a random perturbation and follows Brownian motion biased to return to **0**: at each step, *d***f** /*d*t is sampled from a Gaussian distribution 𝒩 ([− *c*, 0]^*T*^ **f**, **Σ**) which tends to return to **0** at a rate of *c* = 1 and a standard deviation of **Σ** = 0.2**I**. For simplicity, we zeroed the diffusion terms *ν* ^2^∇ **u** and *α* ∇ ^2^s and relied on numerical diffusion (i.e., artificial diffusion due to discretized space and time, numerical instabilities, etc.). This produced a qualitatively acceptable result at higher throughput. We simulated the velocity field at a resolution of 1 mm and the plume intensity at a resolution of 0.5 mm. The time step of the plume simulation is 1/140 seconds. The user can easily modify the parameters of the plume simulation as desired. The simulation is implemented in PhiFlow [42] and based on existing open-source software [43]. Once simulated, we provided the plume concentration s at the appropriate location to the odor sensors of the simulated fly. The plume is also overlaid on the rendered image when applicable.

Then, we implemented a tracking algorithm similar to one presented in a prior study [3] wherein the fly makes decisions based on discrete, binarized plume encounters. While walking, the fly makes turns in a Poisson process with a fixed rate *λ*_turn_. The turn direction is biased toward upwind turns as the encounter frequency increases. This is modeled using a sigmoid function:

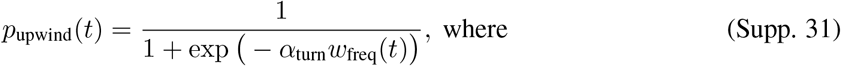

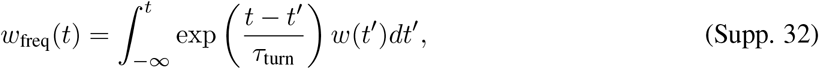

and *p*_upwind_(*t*) is the upwind turn probability, the constant *α*_turn_ = 0.8 Hz^−1^ defines the strength of the upwind bias, *w*_freq_(*t*) is the encounter frequency and is calculated by convolving the encounter history *w*(*t*) with an exponential kernel parameterized by *τ*_turn_ = 2 s. The fly always turns in place for a fixed amount of time t_turn_ = 0.25 s. This leads to similar but variable angles of turning, as the turn might be initiated at different CPG states.

The fly also stops in a Poisson process. The walking-to-stopping Poisson rate *λ*_*w*→*s*_ is reset to a value ∆*λ*_*w*→*s*_ below the baseline *λ*_*w*→*s*,0_ every time the fly encounters the plume, and increases exponentially at a time scale of *τ*_*w*→*s*_ toward the baseline until the next encounter:

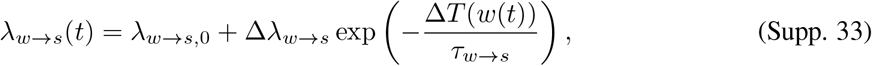

where ∆*T* (*w*(*t*)) is the time since the last encounter. In our simulation, *λ*_*w*→*s*,0_ = 0.78 s^−1^, ∆*λ*_*w*→*s*_ = *−*0.8 s^−1^, τ_*w*→*s*_ = 0.2 s.

Finally, the fly transitions from stopping to walking in a Poisson process as well. Unlike the walking-to-stopping transition, here the Poisson rate *λ*_*s*→*w*_ is based on evidence accumulation rather than a single encounter:

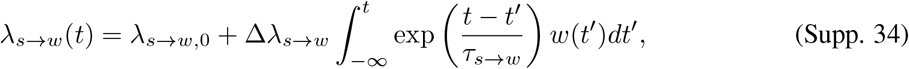

where *λ*_*s*→*w*,0_, ∆*λ*_*s*→*w*_, and *τ*_*s*→*w*_ similarly define the baseline Poisson rate, the amount of change from the baseline, and the time scale of change, respectively. In our simulation, *λ*_*s*→*w*,0_ = 0.5 s^−1^, ∆*λ*_*s*→*w*_ = 1 s^−1^, *τ*_*s*→*w*_ = 0.52 s.

## 2 Supplementary Tables

**Table 1:** Statistics on the locomotor controller benchmark. The U-statistic and p-value in the one-sided, asymptotic Mann–Whitney *U* test used to generate the statistics. “Displacement”: displacement in the forward direction over 1 s. *N* = 20 for each combination of controller and terrain type.

| Displacement |  | CPG < hybrid | Rule-based < hybrid |
| --- | --- | --- | --- |
| | Flat terrain | $U = 346, p = 1.0 \times 10^0$ | $U = 0, p = 3.4 \times 10^{-8}$ |
| | Gapped terrain | $U = 25, p = 1.2 \times 10^{-6}$ | $U = 16, p = 3.5 \times 10^{-7}$ |
| | Blocks terrain | $U = 16, p = 3.5 \times 10^{-7}$ | $U = 34, p = 3.8 \times 10^{-6}$ |
| | Mixed terrain | $U = 43, p = 1.2 \times 10^{-5}$ | $U = 56, p = 5.2 \times 10^{-5}$ |

**Table 2:** Parameters of the CPG-based locomotor controller.

| Variable | Value |
| --- | --- |
| $\phi_{ij}$ | See Table 3, row $i$, column $j$ |
| $w_{ij}$ | 10 if $i \neq j$, otherwise 0 |
| $\nu_i$ | 12 Hz, unless modulated for turning (see “Control of turning”) |
| $R_i$ | 1, unless modulated for turning (see “Control of turning”) |
| $\alpha_i$ | 20 |

**Table 3:**
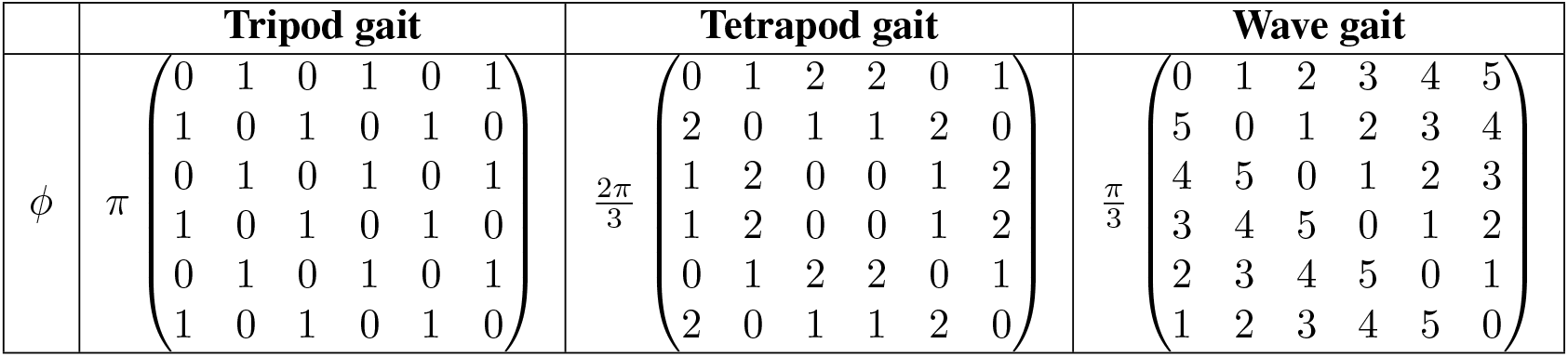
Phase biases *ϕ* between legs for different gaits. The order of rows and columns is right front, right middle, right hind, left front, left middle, and left hind legs.

**Table 4:** Weights used in the rule-based locomotor controller.

|  | Weight |
| --- | --- |
| **Rule 1** | −10 if in swing |
| **Rule 2** (ipsilateral) | $2.5 \text{ s}^{-1}$, cumulative |
| **Rule 2** (contralateral) | $1 \text{ s}^{-1}$, cumulative |
| **Rule 3** (ipsilateral) | $3 \text{ s}^{-1}$, cumulative |
| **Rule 3** (contralateral) | $2 \text{ s}^{-1}$, cumulative |

**Table 5:** Parameters of mechanosensory-based joint angle adjustment rules. Joint angles are adjusted by the specified increment at the specified intervals during leg adjustment and recovery. Note that the roll degree of freedom increment is inverted for the right middle leg.

|  | Degree of freedom | Increment per adjustment | Adjustment interval |  |  |  |
| --- | --- | --- | --- | --- | --- | --- |
|  |  |  | Overstretch rule |  | Stumbling rule |  |
|  |  |  | Lifting | Recovery | Lifting | Recovery |
| **Front leg** | Thorax-coxa pitch | −0.03 rad. | 1.25 ms | 1.43 ms | 0.45 ms | 0.48 ms |
|  | Coxa-trochanter pitch | −0.03 rad. |  |  |  |  |
|  | Femur-tibia | 0.03 rad. |  |  |  |  |
|  | Tibia-tarsus | 0.03 rad. |  |  |  |  |
| **Middle leg** | Thorax-coxa pitch | −0.015 rad. |  |  |  |  |
|  | Thorax-coxa roll | 0.001 rad. |  |  |  |  |
|  | Throax-coxa yaw | 0.025 rad. |  |  |  |  |
|  | Coxa-trochanter pitch | −0.02 rad. |  |  |  |  |
|  | Femur-tibia | −0.02 rad. |  |  |  |  |
| **Hind leg** | Coxa-trochanter pitch | −0.02 rad. |  |  |  |  |
|  | Femur-tibia | 0.01 rad. |  |  |  |  |
|  | Tibia-tarsus | −0.02 rad. |  |  |  |  |

